# Strategic targeting of Cas9 nickase expands tandem gene arrays

**DOI:** 10.1101/2024.09.10.612242

**Authors:** Hiroaki Takesue, Satoshi Okada, Goro Doi, Yuki Sugiyama, Emiko Kusumoto, Takashi Ito

## Abstract

Expanding tandem gene arrays facilitates adaptation through dosage effects and gene family formation via sequence diversification. However, experimental induction of such expansions remains challenging. Here we introduce a method termed break-induced replication (BIR)-mediated tandem repeat expansion (BITREx) to address this challenge. BITREx strategically places Cas9 nickase adjacent to a tandem gene array to break the replication fork that has replicated the array, forming a single-end double-strand break. This break is subsequently end-resected to become single-stranded. Since there is no repeat unit downstream of the break, the single-stranded DNA often invades an upstream unit to initiate ectopic BIR, resulting in array expansion. BITREx has successfully expanded gene arrays in budding yeast, with the *CUP1* array reaching ∼1 Mb. Furthermore, appropriate splint DNA allows BITREx to generate tandem arrays de novo from single-copy genes. We have also demonstrated BITREx in mammalian cells. Therefore, BITREx will find various unique applications in genome engineering.

## INTRODUCTION

Gene duplication plays a critical role in adaptation and evolution, as has become increasingly apparent with the advent of comparative and personal genomics. Two major mechanisms of gene duplication are retrotransposition and non-allelic homologous recombination.^1^ The latter mechanism initially generates a tandemly duplicated pair of the gene, and this configuration often invites additional recombination events to form a tandem gene array. Expansion and contraction of such arrays result in copy number variation (CNV) of the gene.

An immediate effect of expanding tandem gene arrays would be dosage effects. Increased gene dosage often contributes to adaptation to environmental changes. For example, the copy number of an amylase gene, *AMY2B,* shows a notable difference between domestic dogs and wolves, both of which belong to the same species, *Canis lupus*: domestic dogs have more copies than wolves.^2^ This is probably because domestic dogs have adapted to the carbohydrate-rich diet provided by humans, whereas wolves have maintained the ancestral copy number of *AMY2B* without finding any benefit from its high dosage in the wild.

Similarly, the budding yeast *Saccharomyces cerevisiae* shows CNV of the *CUP1* gene, which encodes a copper metallothionein. Different strains have *CUP1* arrays with varying numbers of repeat units, resulting in different levels of copper resistance. Interestingly, the repeat units composing the *CUP1* arrays are not identical across strains; they have distinct breakpoints, suggesting that the initial duplication events from a single copy to two copies occurred independently in their ancestors.^3^ Thus, the formation of the *CUP1* array is an example of convergent evolution. It is considered to have occurred during domestication because *CUP1* is a single-copy gene in many wild isolates of *S. cerevisiae* and in *S. paradoxus*, a non-domesticated cousin of baker’s yeast.^4^ These yeasts are unlikely to have been exposed to high concentrations of copper in the wild, and thus would not have seen any fitness benefit from increased *CUP1* dosage. Expanding tandem gene arrays is a powerful strategy for rapid adaptation to environmental change. Indeed, clones with expanded *CUP1* arrays can be easily obtained in the laboratory by growing yeast cells in the presence of high copper concentrations. ^5^

The expansion of tandem gene arrays can also have long-term effects, including sequence diversification that generates paralogs. This process allows the original tandem gene array to evolve into a genomic locus encoding members of a multigene family. A paradigmatic example of such a locus is the human β-globin locus, which is composed of five genes and one pseudogene. This arrangement allows for the developmental stage-dependent production of three different isoforms—embryonic, fetal, and adult hemoglobins—enabling adaptation to change in oxygen concentration. Extreme examples of gene families with tandemly arrayed paralogs include those for the olfactory receptor (OR), immunoglobulin, and cytochrome P450. For instance, the human and elephant genomes contain ∼400 and ∼2,000 OR genes, respectively, with nearly as many pseudogenes.^6^ While pseudogenization appears to be an inevitable consequence of the functional differentiation of duplicated/multiplicated genes, certain pseudogenes participate in gene conversion, contributing to the immune system, and others play biological roles by generating functional non-coding RNAs.^7^ Tandem gene amplification followed by sequence diversification is therefore a fundamental strategy in evolution by gene duplication.

The recent advent of genome editing has made it possible to manipulate tandem gene arrays. Cleaving the repeat units of a tandem gene array easily induces its contraction through single-strand annealing (SSA) or nonhomologous end joining (NHEJ). In contrast, there is currently no method to expand a tandem gene array. Developing such a method could induce dosage effects and serve as a potential strategy for increasing the yield of useful gene products. It could also provide a basis for generating multigene families, serving as a unique tool for experimental evolution to enhance the potential of cells. But how can we expand a tandem gene array?

In this context, it is interesting to note our previous finding that targeting the catalytically inactive variant of Cas9 (dCas9) to the *CUP1* array induces its contraction in the majority of cells and expansion in a minority of cells.^8^ Mechanistically, dCas9 interferes with replication fork progression, and some of the stalled forks likely break, leading to recombinational repair events that inevitably include non-allelic recombination, resulting in *CUP1* CNV. On the other hand, a single-molecule observation study revealed that replisome disassembles upon collision with Cas9 nickase (nCas9).^9^ Based on our considerations of the potential mechanism for dCas9-induced *CUP1* array expansion and the nCas9-induced replisome disassembly observed by others, we conceived the idea of repurposing break-induced replication (BIR)^10,11^—a mechanism to repair single-end double-strand breaks (seDSBs) generated upon replication fork breakage— to expand tandem gene arrays by strategically targeting nCas9.

## DESIGN

Breakage of a replication fork results in the formation of a seDSB. Subsequent end-resection of the seDSB converts it into a 3′-protruding single-stranded DNA (ssDNA). The ssDNA typically invades the sister chromatid at its allelic position and initiates displacement synthesis, known as BIR, which continues until it encounters a converging replication fork. ^10–12^ When a replication fork collapses within a tandem gene array, the cell can initiate BIR either orthotopically (i.e., at the allelic position) or ectopically (i.e., at non-allelic positions). While orthotopic BIR preserves the tandem gene array, ectopic or out-of-register BIR induces copy number alterations (CNA) of repeat units. If the ssDNA enters a repeat unit downstream or upstream of the seDSB with respect to the direction of replication fork progression, subsequent BIR will decrease or increase the copy number of the repeat units, resulting in contraction or expansion of the array, respectively.

We note that if a replication fork collapses just before completing the replication of a tandem gene array to generate a seDSB within the terminal repeat unit, then the ssDNA derived from the seDSB must invade either the allelic repeat unit or an upstream repeat unit, as there is no downstream repeat unit (Figure 1A). In other words, the array loses the opportunity to contract and thus either remains unchanged or expands. Although we can use nCas9 to induce a seDSB in a replication-dependent manner, we cannot target it selectively to the terminal unit because all repeat units share an identical sequence. However, BIR can also occur, albeit with reduced efficiency, when the invading ssDNA has a 3′-tail sequence that is not homologous to the donor sequence.^13^ Therefore, we hypothesized that targeting nCas9 to the flanking site of a tandem gene array could induce its expansion. We termed this strategy BIR-mediated tandem repeat expansion (BITREx). We first investigated the feasibility of BITREx using the budding yeast *S. cerevisiae* as a model system.

**Figure 1:**
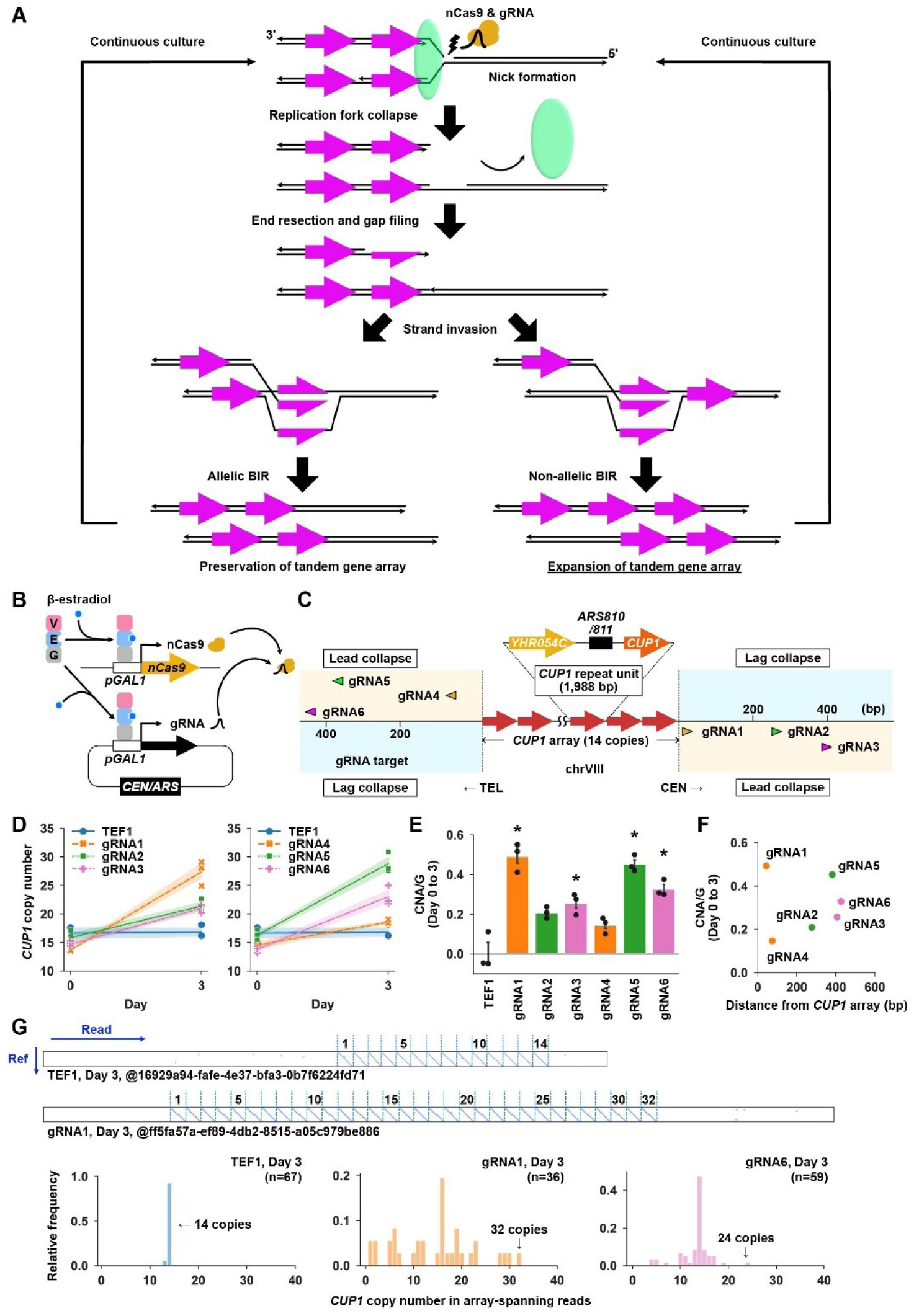
Proof of concept for BITREx by nCas9-induced *CUP1* array expansion. (A) Principle of BITREx. (B) GEV-based system for co-induction of nCas9 and gRNA. (C) *CUP1* array and target sites of effective gRNAs (gRNA1–gRNA6). A rightward or leftward arrowhead indicates that the gRNA sequence is designed for the top or bottom strand, respectively, with its protospacer adjacent motif (PAM). Consequently, the nCas9-gRNA complex nicks the bottom or top strand, which serves as a template for leading strand synthesis initiated from the *ARS810/811* in the *CUP1* repeat unit (lead collapse). (D) nCas9-induced CNA of *CUP1*. Expression of nCas9 and gRNAs were induced by the addition of 10 nM β-estradiol on day 0. Population-averaged *CUP1* copy number was determined by qPCR on days 0 and 3. Each point in the line plot indicates the average *CUP1* copy number (n = 3 biological replicates), with shading around each line indicating the standard deviation (SD). (E) nCas9-induced CNA/G of *CUP1*. CNA was divided by the number of cell divisions estimated from the increase in the optical density of the culture. Error bar represents SD (n = 3 biological replicates). *P < 0.05 (Student’s *t*-test). (F) Plot of the CNA/G value versus the distance from the *CUP1* array to each gRNA target sequence. (G) Nanopore sequencing of *CUP1* arrays. Upper panel, representative dot plots comparing nanopore reads to the *CUP1RU* reference sequence. Lower panel, distribution of *CUP1RU* copy numbers in the specified number of nanopore reads spanning the entire array.

## RESULTS

### Proof of concept for BITREx by nCas9-induced *CUP1* array expansion

We constructed a strain in which β-estradiol induces the expression of Cas9^D10A^, the nCas9 that selectively cleaves the target strand hybridized with guide RNA (gRNA). The induction is mediated by the artificial transcription factor GEV, which consists of the Gal4 DNA-binding domain, estrogen receptor, and VP16 transcriptional activation domain.^14^ Upon binding to β-estradiol, the GEV migrates to the nucleus and activates *GAL1* promoters to induce the expression of Cas9^D10A^ from the genome and its gRNA from a plasmid (Figure 1B). The *CUP1* array is composed of tandem iteration of a ∼2.0-kb DNA segment containing the *CUP1* gene (*CUP1* repeat unit; *CUP1RU*) on chromosome VIII.^3^ It consists mainly of 14 repeat units in the parental strain used in this study (14×CUP1RU, Figures S1A and S1B). We constructed a series of strains that expresses Cas9^D10A^ targeted to the upstream and downstream-flanking sites of *CUP1* array (Figure 1C) and a control site in the *TEF1* locus on chromosome XVI.

We grew these strains by daily dilution of the culture with fresh medium, extracted genomic DNA before and three days after β-estradiol addition, and measured the population-averaged *CUP1* copy number using qPCR (Figure 1D). There was no change in the *CUP1* copy number in the control strain (Figure 1D). In contrast, an increase in the *CUP1* copy number was evident in six out of the 24 strains with Cas9^D10A^ targeted to the flanking regions of the *CUP1* array (Figures 1D, S1C, and S1D). To evaluate the performance of each gRNA, we calculated CNA per generation (CNA/G) (Figure 1E). The six effective gRNAs (gRNA1–gRNA6) showed significantly higher CNA/G values than the other 18 gRNAs (gRNAs1–gRNAs18) and the control *TEF1* gRNA (Figures 1E and S1D). Although the CNA/G values varied among the six gRNAs, they did not correlate with the distance from the *CUP1* array to the gRNA target sites (Figure 1F). Notably, all six gRNAs guide Cas9^D10A^ to nick the template DNA strands for leading strand synthesis initiated by the replication origin *ARS810/811* in the *CUP1RU* (lead collapse).

We intended to investigate the necessity of lead collapse for BITREx. First, we constructed a strain in which the target site of gRNA1, an effective gRNA, is inverted (Figure S1E). In this strain, gRNA1 should make Cas9^D10A^ induce a nick on the lagging strand template. Interestingly, gRNA1 failed to increase *CUP1* copy number in the inverted strain, despite demonstrating comparable cleavage efficiency between the parental and inverted strains when combined with Cas9 (Figure S1F). Next, we intended to investigate Cas9^H840A^, which cleaves the non-target strand not hybridized to the gRNA, thus serving as a complementary nickase to Cas9^D10A^. We used Cas9^H840A/N854A^ in this study due to its reported superiority over Cas9^H840A^ in terms of correct nick formation frequency and avoidance of unwanted indels.^15^ When combined with Cas9^H840A/N854A^, the two gRNAs that performed most efficiently with Cas9^D10A^ (gRNA1 and gRNA6) failed to increase *CUP1* copy number (Figures S2A and S2B). Conversely, two gRNAs that were ineffective with Cas9^D10A^ (gRNAs10 and gRNAs11) induced a small but significant *CUP1* CNA (Figures S2B). In addition, Cas9^H840A/N854A^ with gRNA1 induced a weak increase in the copy number in the inverted strain (Figure S2C). These findings confirm the hypothesized requirement for lead collapse in BITREx. Given the superior efficiency of Cas9^D10A^ over Cas9^H840A/N854A^, we utilized the former nickase throughout this study and will hereinafter refer to it as nCas9 for brevity.

Next, we performed nanopore sequencing of genomic DNA extracted from the strains after three days of BITREx induction. We selected reads containing both 5′- and 3′-flanking regions of the *CUP1* array and generated dot plots between these reads and the *CUP1RU* reference sequence. The number of diagonal lines in the dot plot indicates the number of repeat units comprising the array (Figure 1G). These analyses identified arrays consisting of ∼14, up to 32, and up to 24 copies of *CUP1RU* from strains with *TEF1* gRNA, gRNA1, and gRNA6, respectively (Figure 1G). Note that the apparent discrepancy between the nanopore-based and qPCR-based estimates of *CUP1* copy number arises because the former used only reads that span the entire array, resulting in longer arrays being included less frequently, whereas the latter used all arrays evenly, regardless of their lengths. Intriguingly, nanopore sequencing also revealed the presence of contracted arrays, even though the population-averaged *CUP1* copy number increased (see Discussion for potential causes of the contraction).

We also collected all *CUP1RU*-containing reads and mapped their non-*CUP1RU* sequences to the reference genome. As expected, almost all of them were derived from the *CUP1* array-flanking regions on chromosome VIII (Figure S2D). Since the few abnormal junctions detected were unique to each other, they were most likely artifacts or chimeric molecules generated during the ligation-based library preparation step. The average read depth of chromosome VIII was comparable to the genome-wide average, except for the *CUP1* locus and the polymorphic subtelomeric regions (Figure S2E). Therefore, we concluded that BITREx did not induce detectable levels of translocation involving the *CUP1* locus, ectopic integration of circular DNAs excised from the *CUP1* array, and aneuploidy involving chromosome VIII.

Taken together, targeting nCas9 adjacent to the *CUP1* array induced its expansion *in situ*, proving the principle of BITREx.

### Genetic evidence for BIR to mediate BITREx

To genetically confirm the mechanistic basis of the nCas9-induced *CUP1* array expansion, we constructed a series of strains defective in genes involved in BIR.^10,11^ The initial step of BIR is the invasion of ssDNA into the donor sequence. Rad51 mediates this step in most BIR events. The invaded ssDNA primes the displacement synthesis catalyzed by DNA polymerase δ (Pol δ). Importantly, this step is dependent on Pol32, a Pol δ subunit that is not required for normal S-phase replication. Another essential gene for BIR is *PIF1*, which encodes the DNA helicase indispensable for BIR fork progression. In addition, BIR is suppressed by Rtt109-catalyzed acetylation at Lys-56 of histone H3 (H3K56ac).^16^ Therefore, we tested how the defects of these four genes affect the nCas9-induced increase in *CUP1* copy number using two effective gRNAs (gRNA1 and gRNA6).

First, we found that nCas9 failed to increase *CUP1* copy number in *rad51*Δ and *pol32*Δ strains (Figure 2A) and that episomal expression of *RAD51* and *POL32* suppressed these defects (Figure 2B). Next, we examined the involvement of Pif1 using two mutant alleles, *pif1-m1* and *pif1-m2*.^17^ *PIF1* has two initiation Met codons, the first and second of which direct the synthesis of its mitochondrial and nuclear isoforms, respectively. The *pif1-m1* and *pif1-m2* strains lack the first and second initiation Met codons to synthesize only the nuclear and mitochondrial isoforms, respectively. The *pif1-m2* strain lacking the nuclear isoform was defective in the nCas9-induced *CUP1* copy number increase (Figure 2A), and an episomal *pif1-m1* allele encoding the nuclear isoform suppressed the defect (Figure 2B). Finally, we examined the effect of H3K56ac using *rtt109*Δ and *rtt109 K290Q* alleles (Figure 2C). Rtt109 is the sole enzyme responsible for H3K56ac, but it also contributes to H3K9ac. While no amino acid substitution was known to selectively abolish the H3K56 acetylase activity, K290Q substitution selectively abolishes the H3K9 acetylase activity.^18^ We found that the rate of *CUP1* copy number gain was remarkably enhanced in *rtt109*Δ but not *rtt109 K290Q* cells (Figure 2C). Conversely, the *rtt109 K290Q* allele was as effective as the wild-type *RTT109* allele in suppressing the elevated CNA/G in *rtt109*Δ cells to the wild-type level (Figure 2D). It thus appears that H3K56ac has a negative effect on the nCas9-induced *CUP1* copy number gain. The accelerating effect of *rtt109*Δ on the copy number gain disappeared in the absence of Pol32 (Figure 2C). All these results were consistent with the involvement of BIR in BITREx.

**Figure 2:**
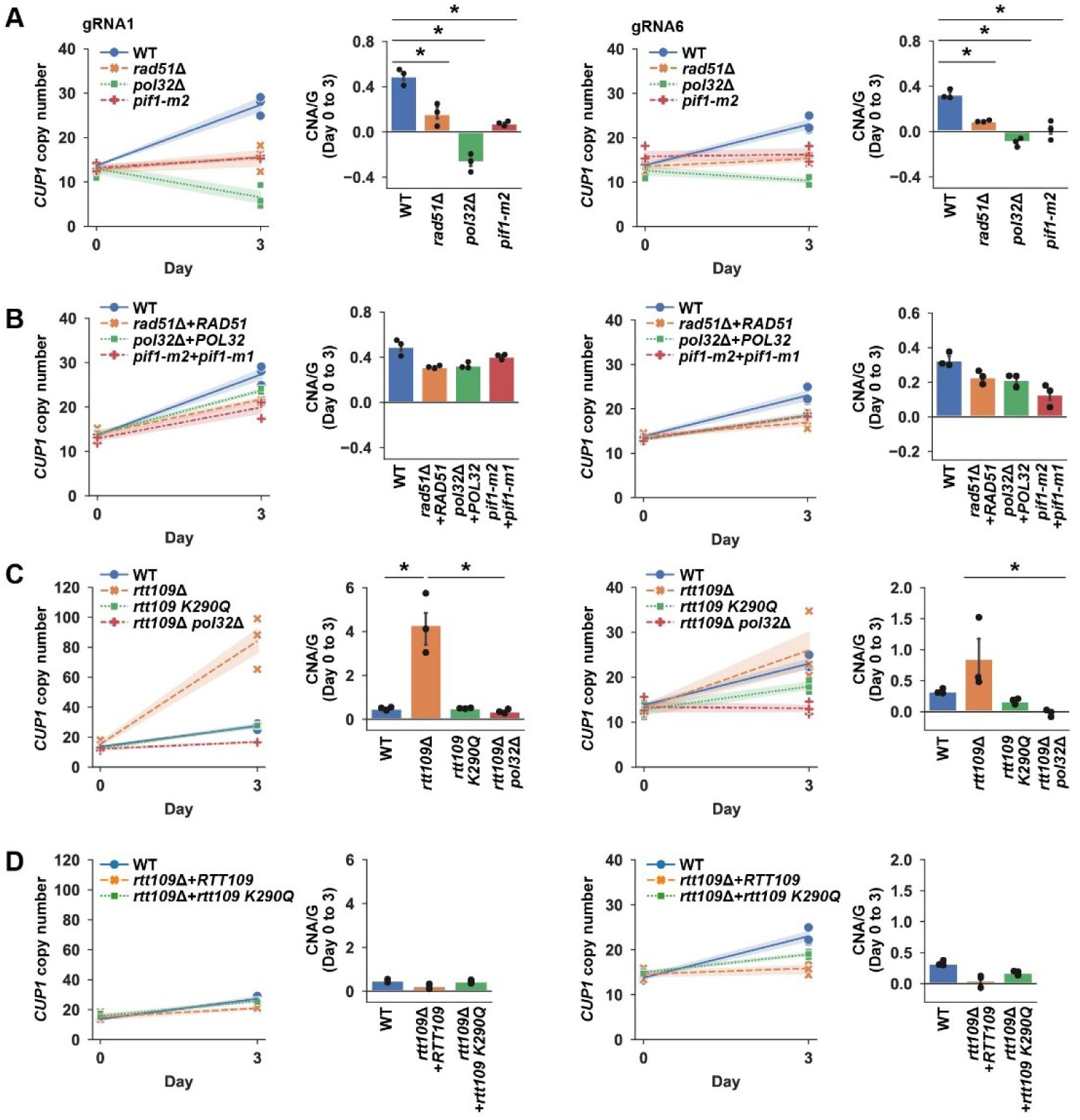
Genetic evidence for BIR to mediate BITREx. (A) BITREx defects in *rad51*Δ, *pol32*Δ, and *pif1-m2* cells. The *CUP1* copy number was measured by qPCR on days 0 and 3 to calculate CNA/G in each mutant. Two gRNAs (gRNA1 and gRNA6) were used. Shading and error bar, SD (n = 3 biological replicates). *P < 0.05 (Student’s *t*-test). (B) Suppression of BITREx defects in *rad51*Δ, *pol32*Δ, and *pif1-m2* cells by episomal copies of *RAD51*, *POL32*, and *pif1-m1*, respectively. Shading and error bar, SD (n = 3 biological replicates). *P < 0.05 (Student’s *t*-test). (C) Enhancement of BITREx in *rtt109*Δ cells but not in *rtt109 K290Q* and *rtt109*Δ *pol32*Δ cells. Shading and error bar, SD (n = 3 biological replicates). *P < 0.05 (Student’s *t*-test). (D) Suppression of BITREx enhanced in *rtt109*Δ cells by episomal copies of *RTT109* and *rtt109 K290Q*. Shading and error bar, SD (n = 3 biological replicates). *P < 0.05 (Student’s *t*-test).

### Long-term BITREx to generate megabase-sized *CUP1* arrays

Theoretically, BITREx occurs every cell cycle to continuously extend the target gene array. To test this possibility, we performed a 31-day continuous BITREx experiment by diluting the yeast cell culture every three days for inoculation into fresh medium. Starting from the wild-type strain with 14 copies of *CUP1RU* (14×CUP1RU), long-term BITREX using gRNA1 or gRNA6, but not *TEF1* gRNA, increased the *CUP1* copy number (Figure 3A). From the ten clones randomly isolated from the gRNA6-expressing cell population with an estimated copy number of ∼200 (Figure 3B), we selected one clone estimated to carry 252 units (250×CUP1RU) for further experiments. Following the curing of the gRNA6-expressing plasmid from this clone, we transformed the cells with the gRNA1- or gRNA6-expressing plasmid and subjected the obtained transformants for another cycle of 31-day BITREx. Intriguingly, the copy number appeared to reach a plateau (∼300) during the second cycle with gRNA6 (Figure 3C). We selected a strain estimated to carry 377 units (380×CUP1RU) from the resulting cell population for subsequent experiments (Figure 3D). In the gRNA1-expressing strain, the copy number appeared to reach a plateau but later showed a decline for unknown reasons (Figure 3C).

**Figure 3:**
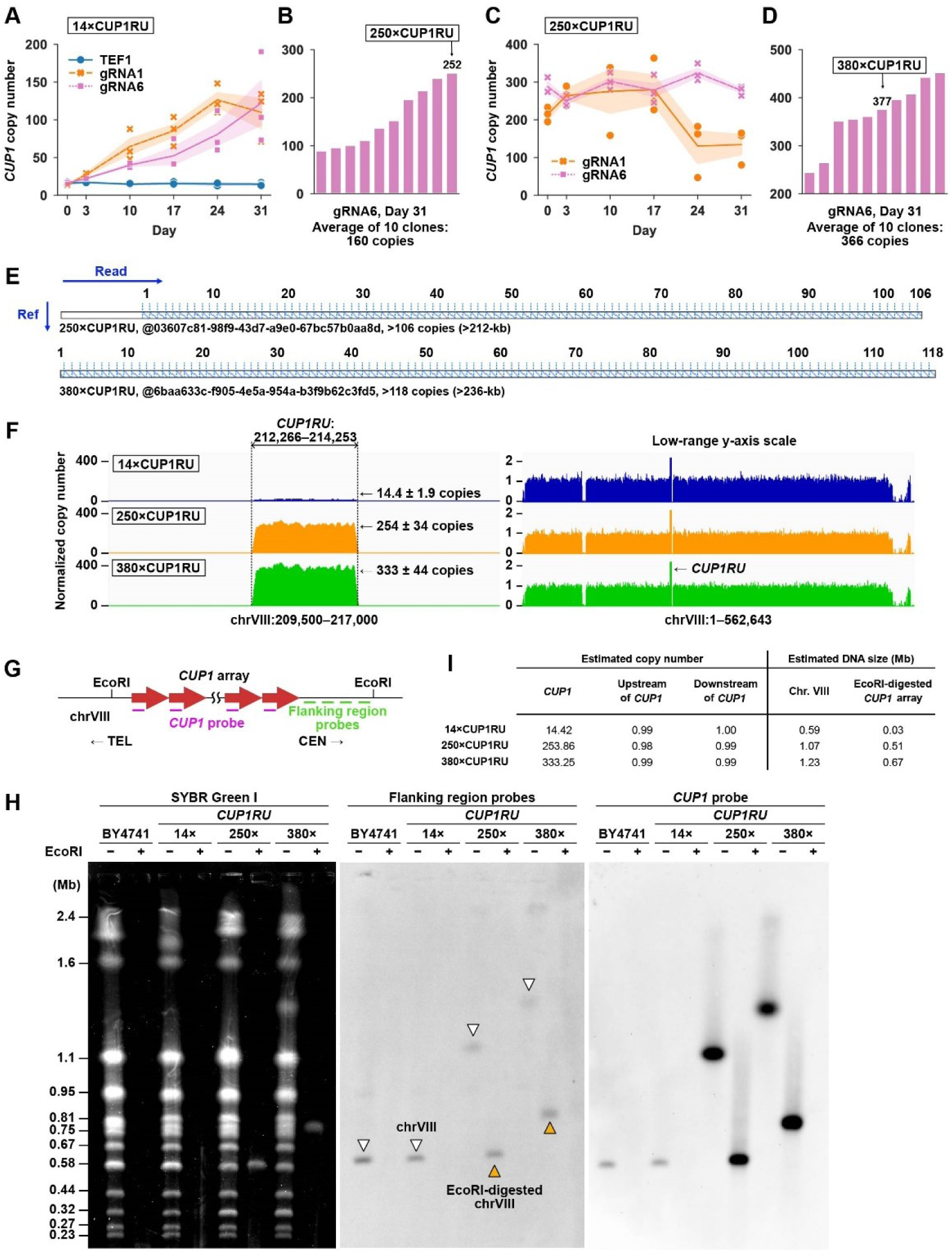
Generation of Mb-sized *CUP1* arrays by long-term BITREx. (A) CNA of *CUP1* over the 31-day BITREx period. Shading, SD (n = 3 biological replicates). (B) *CUP1* copy number of 10 randomly picked clones on day 31 of BITREx using gRNA6. (C) CNA of *CUP1* over the second 31-day BITREx period. Shading, SD (n = 3 biological replicates). (D) *CUP1* copy number of 10 randomly picked clones on day 31 of the second cycle of BITREx using gRNA6. (E) Representative dot plots comparing nanopore reads from the 250×CUP1RU and 380×CUP1RU strains to the *CUP1RU* reference sequence. (F) Normalized read counts in Illumina sequencing. The left panel focuses on the region around *CUP1RU*, while the right panel shows the entire chromosome with a low-range y-axis scale. Read counts were normalized to the average counts of genomic regions excluding rRNA, *CUP1RU*, Ty elements, and mitochondrial DNA. Note that while the sacCer3 reference genome sequence contains two copies of *CUP1RU*, the second copy is masked with ‘N’ prior to mapping. Consequently, the normalized read count directly reflects the *CUP1RU* copy number. The gaps in read counts on the left arm and adjacent to *CUP1RU* are due to the presence of a Ty4 element and the aforementioned masking, respectively. The dip on the right arm results from the presence of a Ty1 element and the segmental duplication between chromosomes VIII and I. (G) Strategy of Southern blot hybridization. (H) PFGE analysis of *CUP1* arrays expanded by long-term BITREx. Left, SYBR Green I stain; middle, blot hybridized with the flanking region probes; right, blot hybridized with the *CUP1* probe. White and orange arrowheads indicate chromosome VIII and the EcoRI fragment containing the *CUP1* array, respectively. (I) Deep sequencing-based estimates of the sizes of chromosome VIII and the EcoRI restriction fragment containing the *CUP1* array.

We performed nanopore sequencing to detect the expanded *CUP1* array in the 250×CUP1RU and 380×CUP1RU strains. Among the reads obtained are those containing more than 106 (>212 kb) and 118 (>236 kb) copies of *CUP1RU* (Figure 3E). However, we failed to obtain such reads that span the full length of the extended array, which must be longer than ∼500 and ∼760 kb for 250×CUP1RU and 380×CUP1R, respectively. Nevertheless, the normalized read counts of *CUP1RU* in Illumina sequencing were 254 ± 34 and 333 ± 44 in the 250×CUP1RU and 380×CUP1RU strains, respectively, while those of non-*CUP1RU* regions on chromosome VIII were equal to one (i.e. genome-wide average) (Figure 3F), suggesting that BITREx can expand the *CUP1* array *in situ* to a sub-Mb size.

To visualize the entire chromosome VIII and the *CUP1* array in these strains, we performed pulsed-field gel electrophoresis (PFGE) followed by Southern blot hybridization using probes derived from either *CUP1RU* or its flanking region (Figure 3G). Both probes hybridized to 0.6-, 1.1-, and 1.4-Mb bands in the intact genomic DNAs derived from the 14×CUP1RU, 250×CUP1RU, and 380×CUP1RU strains, respectively (Figure 3H). We also examined EcoRI-digested genomic DNA. Since *CUP1RU* has no EcoRI site, the extended arrays appeared as 0.5- and 0.8-Mb bands in the 250×CUP1RU and 380×CUP1RU strains, respectively (Figure 3H). These sizes were consistent with those estimated from the Illumina deep sequencing (Figure 3I). We did not detect any aberrant bands indicating translocations or other chromosomal alterations.

To further explore the potential of BITREx in *CUP1* array expansion, we performed a long-term culture experiment using the *rtt109*Δ strain, as it remarkably accelerated the array expansion (Figure 2C). In the absence of Rtt109, BITREx for 31 days increased the average *CUP1* copy number to >500, corresponding to arrays longer than 1 Mb (Figure S3A). We examined individual colonies by qPCR and obtained strains with estimated copy numbers of 274 (*rtt109*Δ 250×CUP1RU), 387 (*rtt109*Δ 380×CUP1RU), and 819 (*rtt109*Δ 800×CUP1RU) for further experiments (Figure S3A). After three days of culture without estradiol, the first two strains exhibited only a modest decrease in the *CUP1* copy number as did the wild-type strains with similar copy numbers, while the third strain showed a significant decrease from ∼700 to ∼350 copies (Figure S3B). The normalized Illumina read counts in the *CUP1RU* increased significantly, whereas those in the non-*CUP1RU* regions on chromosome VIII did not show any detectable CNAs, ruling out the possibility of aneuploidy (Figure S3C). Although Southern blot hybridization confirmed the elongation of chromosome VIII and the *CUP1* array, the hybridization bands in *rtt109*Δ strains were much broader than those in wild-type strain (Figure S3D). Therefore, *rtt109*Δ cells likely have more heterogeneous *CUP1* arrays than wild-type cells.

Taken together, BITREx can expand the *CUP1* array to Mb size, especially in the absence of Rtt109. To maintain the extremely long arrays, BITREx likely needs to be continuously induced.

### Epigenetic modulation of BITREx

To further investigate the role of Rtt109-generated H3K56ac, we examined the effect of nicotinamide (NAM), which inhibits the NAD^+^-dependent histone deacetylase family consisting of Sir2, Hst1, Hst2, Hst3, and Hst4 in the budding yeast.^19^ Previous studies have reported that NAM induces *CUP1* copy number reduction in an Rtt109/H3K56ac-dependent manner.^8,20^ We first confirmed that H3K56ac accumulates in the 14×CUP1RU strain after 24 h of NAM exposure (Figure 4A). Notably, the presence of NAM not only suppressed BITREx of the *CUP1* array (Figure 4B), but also led to its gradual contraction, regardless of whether nCas9 was targeted to the *CUP1*-flanking site or the control site (Figure 4C). The effect of NAM on highly-extended *CUP1* arrays was remarkable: the 250×CUP1RU and 380×CUP1RU strains showed a drastic decrease in the copy number during three days of NAM exposure (Figure 4D).

**Figure 4:**
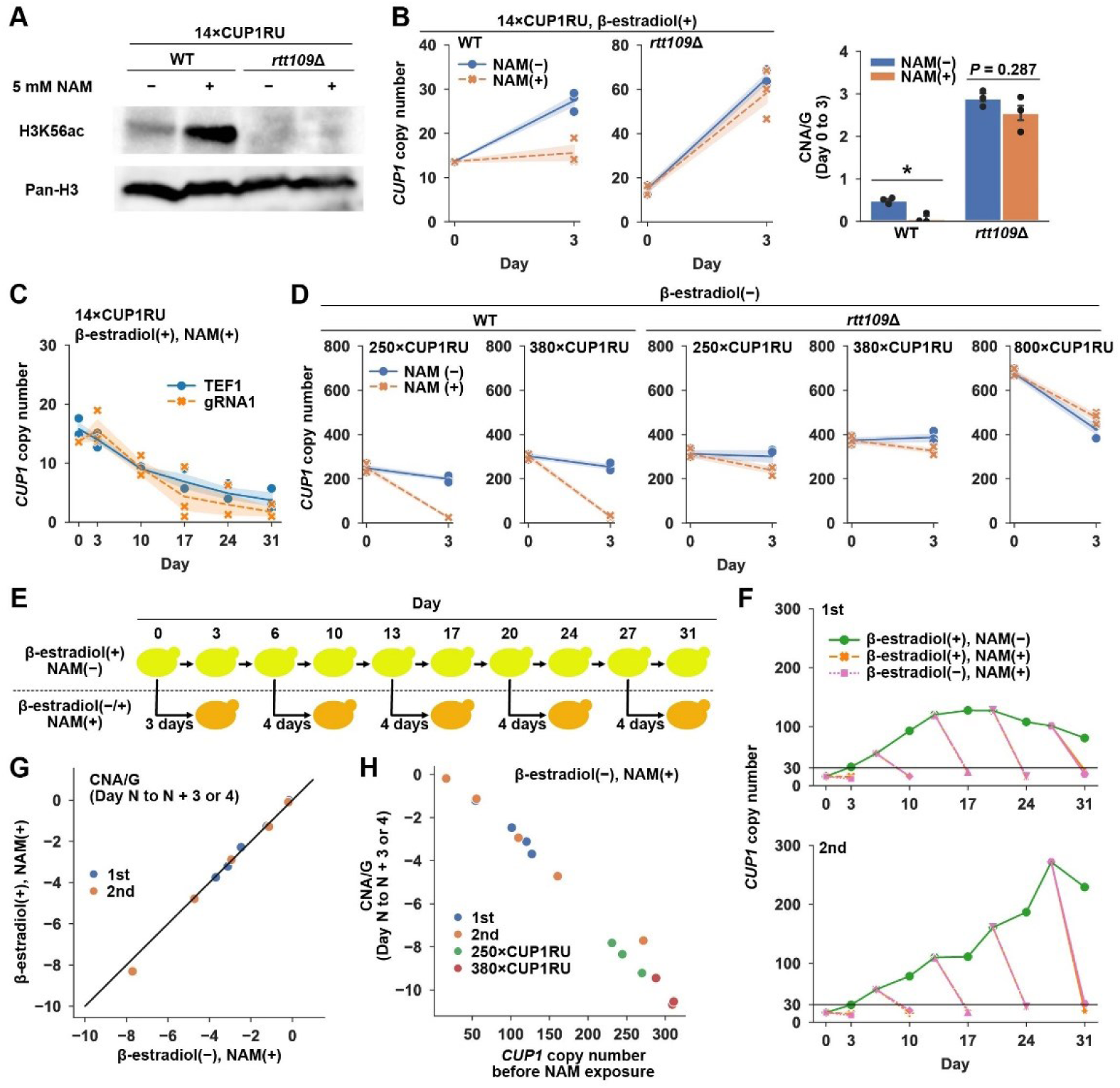
Epigenetic modulation of BITREx. (A) Effects of NAM on H3K56ac. The wild-type and *rtt109*Δ cells were grown in the absence and presence of 5 mM NAM for three days and subjected to immunoblot analysis of H3K56ac and total histone H3. (B) Effects of NAM on BITREx of the *CUP1* array using gRNA1. Shading and error bar, SD (n = 3 biological replicates). *P < 0.05 (Student’s *t*-test). (C) Effects of long-term NAM exposure on the normal *CUP1* array in the wild-type strain (14×CUP1RU) with *TEF1* gRNA and gRNA1. Shading, SD (n = 3 biological replicates). (D) Effects of short-term NAM exposure on the BITREx-extended *CUP1* arrays in the presence and absence of Rtt109. Note that BITREx was not induced during the NAM exposure. Shading, SD (n = 3 biological replicates). (E) Experimental strategy to examine the effect of initial *CUP1* copy number on NAM-induced array contraction. (F) NAM-induced contraction of variably extended *CUP1* arrays. Line plots indicate the actual data following the strategy depicted in (E) (n = 2 biological replicates). BITREx was induced in the wild-type strain using gRNA1. The horizontal lines indicate 30 copies. (G) Effect of BITREx on NAM-induced contraction of extended *CUP1* arrays. The CNA/G values are compared between the absence (*x*-axis) and presence (*y*-axis) of BITREx induction. The diagonal line indicates *y* = *x*. (H) Plot of CNA/G versus initial *CUP1* copy number. Data from (F) and (D) (i.e., 250×CUP1RU and 380×CUP1RU strains) were used.

These results suggested that the effect of NAM on CNA depends on the initial length of the *CUP1* array. To test this hypothesis, we prepared a series of cell populations with different average *CUP1* copy numbers by temporal sampling from a long-term BITREx culture. Each sample was divided into two subpopulations, which were then cultured in parallel in the presence and absence of NAM for three or four days (Figure 4E). In all samples, NAM exposure induced a steep decrease in *CUP1* copy number, converging to <30 copies (Figure 4F). There was no difference in CNA/G levels with or without nCas9 induction (Figure 4G), likely because NAM efficiently suppresses BITREx (Figure 4B). The CNA/G appeared to inversely correlate with the initial *CUP1* copy number (Figure 4H), as expected from the theoretical model, which assumes array contraction via homologous recombination between repeat units following second-order kinetics (see Methods).

Note that the effects of NAM on BITREx and the *CUP1* array were completely abolished in the absence of Rtt109, the sole H3K56 acetylase (Figures 4A, 4B, and 4D). These results indicate that although NAM leads to the accumulation of acetylation at multiple Lys residues, its effects are mediated through the accumulated H3K56ac.

### General applicability of BITREx

Theoretically, BITREx can extend a tandem array consisting of two or more repeat units but not a single-copy unit. We constructed strains harboring *CUP1* arrays consisting of one, two, and three repeat units (i.e., 1×CUP1RU, 2×CUP1RU, and 3×CUP1RU) to determine the minimum number of repeat units required for BITREx (Figure 5A). When targeted to an upstream or downstream flanking site of the *CUP1* array for three days, nCas9 increased the *CUP1* copy number in the 2×CUP1RU and 3×CUP1RU strains, but not in the 1×CUP1RU strain (Figure 5B). Nanopore sequencing revealed the expanded *CUP1* arrays in the 2×CUP1RU and 3×CUP1RU strains but not in the 1×CUP1RU strain (Figure 5C). All these results are consistent with the BIR-based mechanism of BITREx.

**Figure 5:**
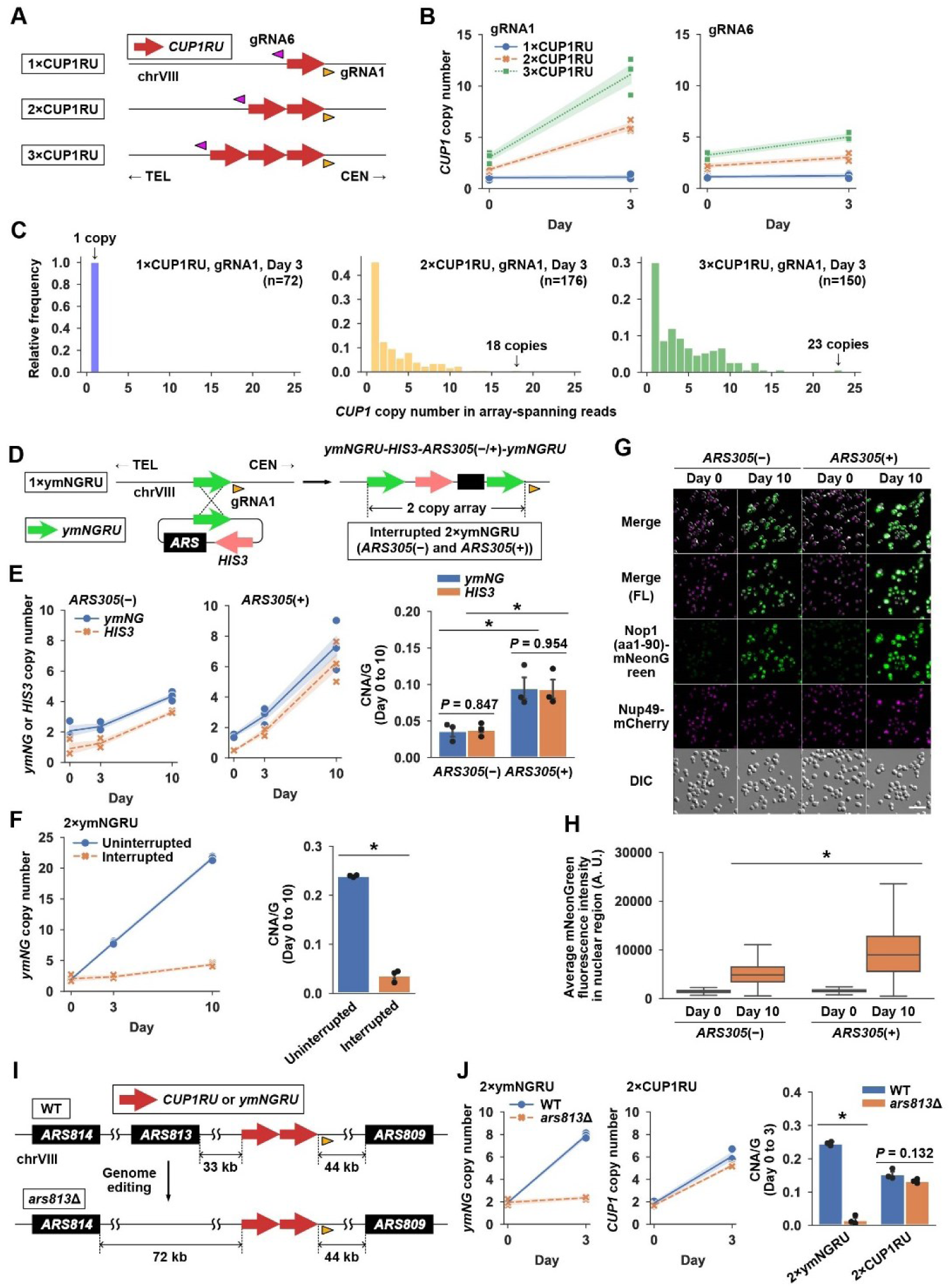
Core requirements and modulating factors for BITREx. (A) Schematic of the strains with one to three copies of *CUP1RU*. Orange and magenta arrowheads indicate the target sites of gRNA1 and gRNA6, respectively. (B) *CUP1* CNA in the three strains depicted in (A). Shading, SD (n = 3 biological replicates). (C) Distribution of *CUP1* copy number in nanopore reads spanning the entire array. Nanopore sequencing was performed on day 3 of BITREx. (D) Plasmid integration strategy to generate an interrupted *2×ymNGRU* array. The intervening sequence was composed of the plasmid backbone containing *HIS3* with or without *ARS305*. (E) CNA of *ymNGRU* and *HIS3* in the absence and presence of the embedded *ARS305*. Shading and error bar, SD (n = 3 biological replicates). *P < 0.05 (Student’s *t*-test). (F) CNA of *ymNGRU* in the uninterrupted and interrupted two-unit arrays. Shading and error bar, SD (n = 3 biological replicates). *P < 0.05 (Student’s *t*-test). (G) Microscopic images of strains bearing the interrupted *2×ymNGRU* arrays without and with the embedded *ARS305*. These strains have *NUP49-mCherry* to visualize the nuclei (magenta). FL, fluorescence; DIC, differential interference contrast. (H) Quantification of mNeonGreen fluorescence in (G). Box plots indicate the distribution of the average mNeonGreen fluorescence intensity in the nuclear region. The bottom and top of the box show the first and third quartiles, respectively. The bar in each box represents the median value, and the error bars represent the range of values. *P < 0.001 (one-way ANOVA test). (I) ARS distribution around the *CUP1* locus in the wild-type and *ars813*Δ strain. Arrowhead, gRNA1-target site. (J) Effect of *ARS813* on BITREx of the uninterrupted *2×ymNGRU* and *2×CUP1RU* arrays. The *2×ymNGRU* array lacks internal ARS, while the *2×CUP1RU* array contains *ARS810/811* within the repeat unit. Shading and error bar, SD (n = 3 biological replicates). *P < 0.05 (Student’s *t*-test).

Since all the results described so far were on the *CUP1* array, we next intended to test whether BITREx is applicable to other natural tandem gene arrays. We focused on the *ENA1/2/5* array encoding P-type ATPase sodium pumps.^21^ The *ENA1/2/5* array comprises a tandem array of three paralogous genes, namely *ENA1*, *ENA2*, and *ENA5*, on chromosome IV in the S288C reference genome sequence. However, other strains were reported to have four or more paralogs,^22,23^ and the strain used in this study has five paralogs (Figure S4A). We succeeded in BITREx of the *ENA1/2/5* array (Figures S4B–S4E).

We also tested whether BITREx could expand engineered or synthetic gene arrays. To investigate this, we designed a ∼1.6-kb repeat unit called *ymNGRU*. This unit consists of a yeast codon-optimized coding sequence for the fluorescent protein mNeonGreen (*ymNG*), preceded by the *NOP1* promoter and the coding sequence for the nuclear localization signal-containing domain of Nop1 (amino acid residues 1–90), and followed by the *ADH1* terminator (Figure S4F). We successfully expanded a two-unit array of *ymNGRU* (*2×ymNGRU*) that replaced the *CUP1* array in a disruptive manner (Figures S4G–S4I).

Finally, we tested whether BITREx could expand engineered gene array at loci that do not form natural tandem repeats. For this purpose, we inserted a two-unit array of *CUP1RU* or *ymNGRU* with the gRNA1-target sequence into the *HO* locus on chromosome IV (*ho*Δ*::2×CUP1RU* or *ho*Δ*::2×ymNGRU*) (Figure S5A). Note that these arrays include *HIS3* between the two repeat units to facilitate strain construction. They can thus be interpreted as interrupted two-unit arrays of *CUP1RU/ymNGRU*. We inserted the interrupted two-unit array in two orientations: in one strain, the nick can be introduced on the *ARS404*-proximal side, and in the other strain, on the *GCS1*-proximal side of the array (Figure S5A). The replisome originating from *ARS404*, located approximately 50 bp telomeric to the *HO* gene, is likely responsible for the replication of this locus (Figure S5A).^25^ As expected from the principle of BITREx, we were able to expand these arrays much more effectively in the strains with the *GCS1*-proximal nick than in those with the *ARS404-*proximal nick (Figures S5B and S5C), highlighting the importance of understanding the replication fork directionality for efficient BITREx design. We also succeeded in BITREx of these two-unit arrays integrated at the *X-2* locus, an intergenic safe harbor site on chromosome X^26^ (Figures S5D–S5F).

Taken together, these results demonstrate that BITREx can expand two-unit arrays, whether uninterrupted or interrupted, at different genomic loci. However, the efficiency of expansion varies depending on the locus and the composition of the array.

### Effects of ARS on BITREx

The successful expansion of the interrupted two-unit arrays prompted us to explore the possibility of BITREx-mediated amplification of a target sequence embedded between two repeat units, as this configuration can be easily generated using the conventional plasmid integration technique. For example, we integrated a ∼5-kb plasmid carrying *ymNGRU* and *HIS3* into the single-copy *ymNGRU* at the *CUP1* locus (Figure 5D) and successfully applied BITREx to the resulting interrupted two-unit array, leading to a simultaneous increase in the copy numbers of *ymNGRU* and *HIS3* (Figure 5E). However, the interrupted *2×ymNGRU* array showed a significantly lower CNA/G than the uninterrupted *2×ymNGRU* array (Figure 5F). Given the importance of replication fork directionality (Figures S5A–S5C), we were concerned that the ∼3.4-kb plasmid-derived intervening sequence might have decreased the likelihood of achieving the desired fork direction at the nick. To mitigate such adverse effects, we incorporated an autonomously replicating sequence (ARS) into the repeat unit. As expected, incorporating *ARS305* effectively increased CNA/G and enhanced the fluorescence (Figures 5E, 5G, and 5H).

These results led us to hypothesize that BITREx of a tandem gene array without an internal ARS would be sensitive to the distance between the nick on one side and the nearest ARS on the opposite side of the array. In contrast, BITREx of a tandem gene array with an ARS within the repeat unit would not be affected by this distance. To test this hypothesis by elongating the distance between the nick and the ARS, we generated strains deleted for *ARS813*, the nearest ARS responsible for BITREx using gRNA1 at the *CUP1* locus (Figure 5I). Indeed, BITREx of the *2×ymNGRU* array, which lacks an internal ARS, depended on *ARS813*, while BITREx of the *2×CUP1RU* array, which includes *ARS810/811*, did not (Figure 5J). The incorporation of an ARS had no negative effect on BITREx and even enhanced it. Therefore, it is desirable to incorporate an ARS in the repeat unit, especially when no suitable external ARS is available near the target locus or when the repeat unit is long.

Based on these results, we attempted to amplify a multigene array using BITREx. We used plasmid integration to construct strains where the two-unit *CUP1* array at the *CUP1* locus was interrupted by a ∼10-kb fragment containing yeast codon-optimized coding sequences for four fluorescent proteins (mTagBFP, miRFP682, mCherry, and mNeonGreen) and *HIS3*, with and without *ARS305* (Figure S5G). BITREx successfully extended the array, particularly in the strain with *ARS305* in the repeat unit (Figure S5H), enhancing the dosage effects of the four passenger genes or fluorescence (Figure S5I). We also observed that the *HO* and *X-2* loci successfully supported BITREx of the multigene array, especially in the presence of *ARS305*, though with lower CNA/G values compared to the *CUP1* locus (Figure S5H). Intriguingly, the effects of *ARS305* were evident between days 3 and 10 but not between days 0 and 3 (Figure S5H). The CNA/G between days 3 and 10 is significantly lower than that between days 0 and 3 in the absence of *ARS305*, but not in its presence (Figure S5J). This is presumably because the desirable replication fork directionality at the nick was similar between ARS-less and ARS-containing arrays while they remained relatively short, but could not be maintained in the ARS - less arrays as they expanded, unlike in the ARS-containing arrays.

### Splinted BITREx for *de novo* generation of tandem gene arrays

The minimum requirement for BITREx is two identical sequences to form either an uninterrupted or interrupted two-unit array. However, because BIR frequently switches templates,^27^ we hypothesized that BITREx could generate a tandem array starting from a single-copy sequence if an engineered DNA serves as a “splint” to mediate the necessary template switching.

To test this idea, we constructed a strain that harbors a repeat unit delimited by *U2* and *LE* (two non-overlapping consecutive fragments of the *LEU2* coding sequence) on chromosome VIII and a plasmid carrying the splint fragment *EU* (Figure 6A). Upon the induction of a seDSB by targeting nCas9 to a downstream-flanking site of the genomic *LE*, the ssDNA generated by end-resection of the break site may use its *E* to invade the episomal *EU* for further extension by displacement synthesis. The extended ssDNA may then use the newly acquired *U* to switch its template from the splint plasmid to the genomic *U2*. If these consecutive strand invasion-extensions occur, the *U2–LE* unit duplicates to reconstitute the *LEU2* gene, allowing the growth in the absence of leucine. Indeed, we observed the emergence of Leu^+^ cells with different efficiencies depending on the length of the splint fragment (Figure 6B). Note that the nicking of the splint plasmid was critical for the efficiency (Figure 6B).

**Figure 6:**
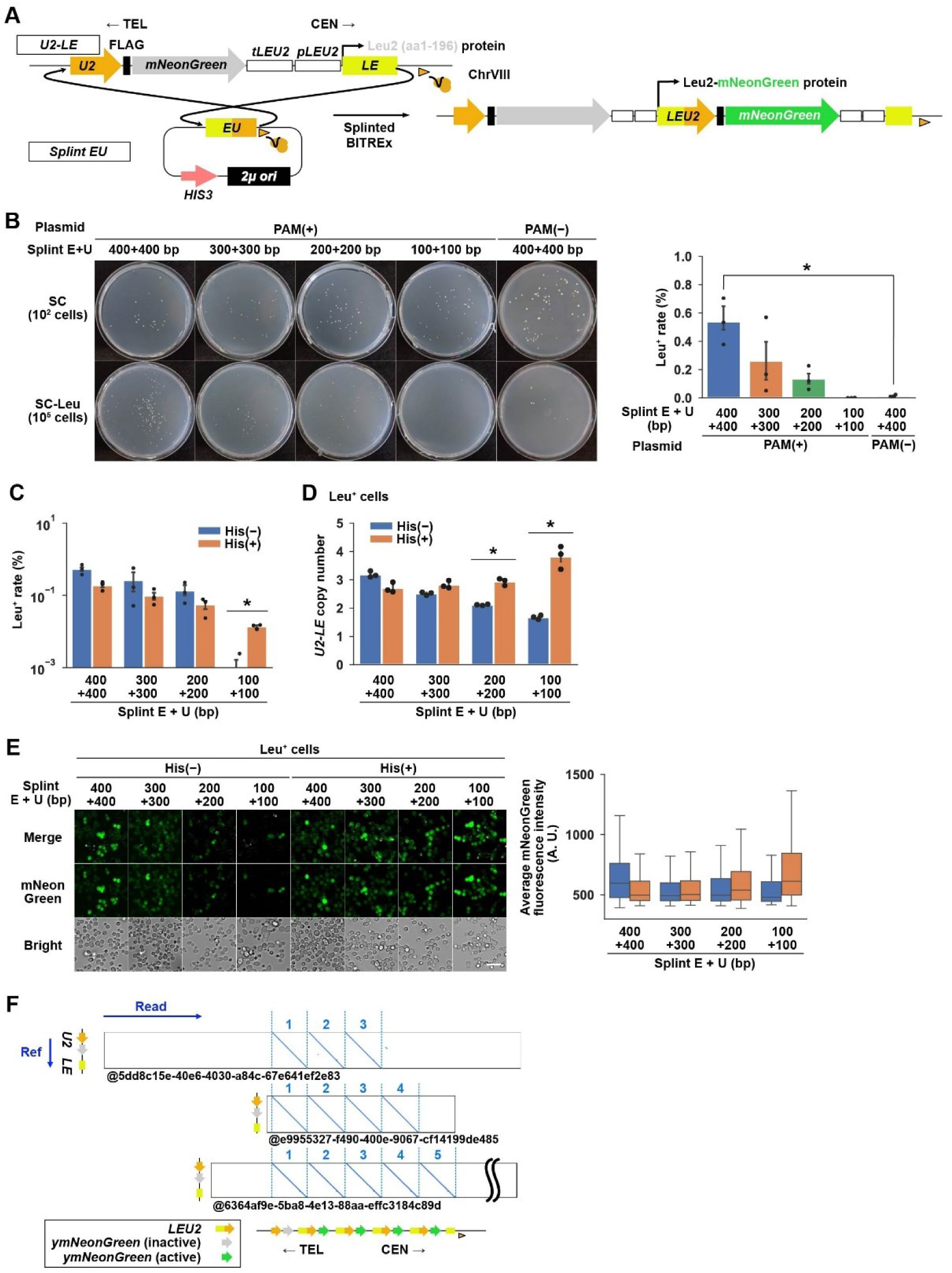
De novo generation of tandem gene array by splinted BITREx. (A) Proof-of-concept experiment for splinted BITREx. A single round of splinted BITREx reconstitutes the *LEU2*-coding sequence, resulting in the expression of a fusion protein composed of Leu2 and mNeonGreen connected by the FLAG tag. *tLEU2*, *LEU2* terminator; *pLEU2, LEU2* promoter; orange arrowhead, gRNA1-target site. (B) Efficiency of splinted BITREx measured by the appearance of Leu^+^ colonies. Splints of four different lengths were used in 3-day BITREx. Representative images are shown for colonies appeared on SC and SC−Leu agar plates. The bar chart displays the fraction of Leu^+^ cells among the living cells. Error bar, SD (n = 3 biological replicates). *P < 0.05 (Student’s *t*-test). (C) Effects of histidine supplementation on the appearance of Leu^+^ colonies. Error bar, SD (n = 3 biological replicates). *P < 0.05 (Student’s *t*-test). (D) Effects of histidine supplementation on the copy number of *U2–LE* unit. Error bar, SD (n = 3 biological replicates). *P < 0.05 (Student’s *t*-test). (E) Effects of histidine supplementation on the mNeonGreen fluorescence. Left, microscopic images of strains subjected to splinted BITREx. Right, box plots showing the distribution of the average mNeonGreen fluorescence intensity in cells. (F) Dot plot comparing nanopore reads to the reference sequence of *U2–LE* unit.

Once the *LEU2* gene is reconstituted, the *LE* ssDNA can initiate BIR either by directly invading the reconstituted *LEU2* or indirectly invading the genomic *U2* or *LEU2* via the splint plasmid (Figure 6A). Since the direct invasion should be much more efficient than the indirect invasion, the splint plasmid may inhibit subsequent cycles of BITREx, even though it is essential for the initial reconstitution of *LEU2*. Based on these considerations, we examined the effect of histidine supplementation starting from day 2 to promote the spontaneous loss of the *HIS3*-marked splint plasmid. This supplementation protocol improved the emergence of Leu^+^ clones when combined with the shortest splint fragments (Figure 6C). It also increased the copy number of *U2–LE* units and the mNeonGreen fluorescence when combined with the shortest or second shortest splint fragment (Figures 6D and 6E). Nanopore sequencing identified reads spanning a tandem array composed of up to five *U2–LE* units or four *LEU2-mNeonGreen*-fusion genes (Figure 6F).

These results demonstrated that “splinted BITREx” enables *de novo* formation of a tandem gene array from a single-copy sequence.

### BITREx in mammalian cells

We next investigated the feasibility of BITREx in mammalian cells. Since BIR has been demonstrated in mammalian cells using the reconstitution of a fluorescent protein gene,^28^ we generated a reporter construct *mCherry-PuroR-FP-SV40ori-EGF-gRNA1 target site* (Figure 7A). In this construct, the *mCherry-PuroR* portion serves as a transfection reporter/marker, while the *FP-SV40ori-EGF* portion serves as an interrupted two-unit array, in which two *F* fragments are interrupted by the *P-SV40ori-EG* fragment. Therefore, nCas9 with gRNA1 should induce BIR via the *F* fragment to reconstitute the *EGFP* gene, resulting in green fluorescence.

**Figure 7:**
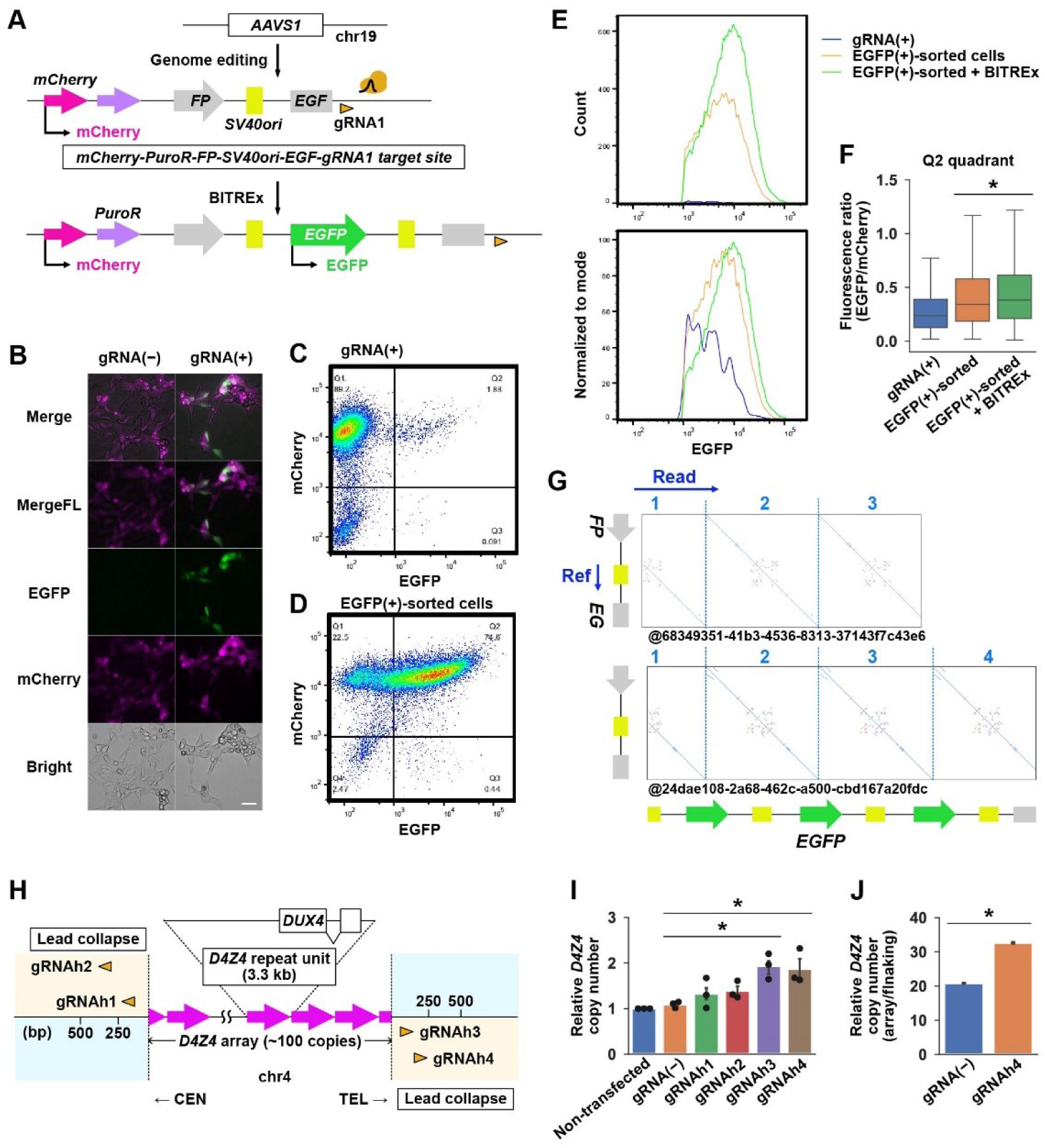
BITREx in mammalian cells. (A) Proof-of-concept experiment for BITREx in HEK293T cells using an *EGFP* reconstitution reporter integrated into the *AAVS1* locus. (B) Microscopic images of HEK293T cells with the integrated reporter on day 4 after transfection of the nCas9-gRNA1 expression plasmid. (C) Flow cytometric analysis of mCherry and EGFP expression on day 3 after transfection of the nCas9-gRNA1 expression plasmid. The cells gated based on light scatter (Figure S7B) displayed mCherry and EGFP expression across four quadrants (Q1, Q2, Q3, Q4), with the percentage distribution indicated in each quadrant. (D) Flow cytometric analysis of mCherry and EGFP expression in the mCherry^+^/EGFP^+^ cells sorted from quadrant Q2 in (C). (E) Distribution of EGFP fluorescence. Blue, the total population of cells transfected with the gRNA1-nCas9 co-expression plasmid; orange, flow-sorted EGFP^+^ cells; green, flow-sorted EGFP^+^ cells on day 3 after re-transfection with the co-expression plasmid. The *x*-axis represents EGFP fluorescence intensity, while the *y*-axis represents either the number of cells (top) or the value normalized to mode (bottom). (F) Box plots showing the fluorescence ratio of EGFP to mCherry in the cells in (E). *P < 0.05 (Student’s t-test). (G) Dot plots comparing nanopore reads to the reference sequence of the *FP–EG* unit. The upper and lower reads indicate at least triplication and quadruplication, respectively. (H) Schematic of the *D4Z4* array and its flanking regions on human chromosome 4. Arrowheads indicate the target sites of four gRNAs, which guide nCas9 to induce lead collapse of the replication fork moving outward from the *D4Z4* array. (I) *D4Z4* copy number quantified by qPCR on day 3 of BITREx with the indicated gRNA. Error bar, SD (n = 3 biological replicates). *P < 0.05 (Student’s *t*-test). (J) Ratio of normalized read counts between the *D4Z4* macrosatellite and the centromeric flanking region.

We integrated the construct to the safe harbor locus *AAVS1* of human HEK293T cells, which expresses the large T antigen that activates *SV40ori*, the replication origin of the SV40 virus, using the VIKING method for efficient NHEJ-based knock-in.^29^ Following the puromycin selection of cells with integrated construct, we transfected the selected cells with a plasmid that co-expresses nCas9 and gRNA1 to induce BITREx (Figure S6A). As a fraction of the transfected cells started to show EGFP fluorescence (Figure 7B), we flow-sorted the mCherry^+^/EGFP^+^ cells (Figures 7C, 7D, and S6B). A subsequent round of transfection with the co-expression plasmid conferred enhanced EGFP fluorescence to the cells (Figure 7E). As expected from the design of the reporter construct, in which BITREx increases the copy number of the *EGFP* gene but not the *mCherry* gene (Figure 7A), the EGFP/mCherry fluorescence ratio increased (Figure 7F). Nanopore sequencing of these cells identified reads containing at least three or four copies of the *FP-SV40ori-EG* unit (Figure 7G). These results demonstrated the feasibility of BITREx in mammalian cells.

We then aimed to apply BITREx to gene-sized tandem repeats naturally occurring in the human genome. For this purpose, we focused on the *D4Z4* array, which consists of a 3.3-kb repeat unit on chromosome 4q35. This array is significant because its heterozygous contraction causes facioscapulohumeral muscular dystrophy (FSHD), the third most common type of inherited muscular dystrophy.^30^ Accordingly, its expansion may have potential implications in FSHD therapeutics. We designed four gRNAs to target nCas9 to the centromeric and telomeric sides of the *D4Z4* array (Figure 7H). The results of qPCR consistently demonstrated that targeting nCas9 to the telomeric side increased the *D4Z4* copy number (Figure 7I). We subjected the cells with no gRNA and the most effective gRNA (gRNAh4) to nanopore sequencing. The normalized read count indicated that the *D4Z4* copy number increased from ∼20 to ∼32 copies (Figure 7J), consistent with the qPCR results (Figure 7I). Although the complexity of the *D4Z4* locus containing many repetitive sequences and the presence of an almost identical locus on chromosome 10q26 precluded the complete characterization of the induced CNAs, these results demonstrated the applicability of BITREx to endogenous tandem gene arrays in mammalian genomes.

## DISCUSSION

We have developed BITREx, a method for expanding tandem gene arrays through continuous ectopic BIR induced by strategically targeting nCas9. In BITREx, nCas9 is placed adjacent to a tandem gene array, disrupting the replication fork after it has replicated the array. We previously developed Paired Nicking-induced Amplification (PNAmp), a method for inducing large segmental duplications by paired nicking.^31^ While both PNAmp and BITREx induce structural variations by manipulating replication fork progression, they are mechanistically distinct: PNAmp uses two gRNAs and depends on SSA,^31^ whereas BITREx uses one gRNA and relies on BIR.

For BITREx to be effective, the replication fork crossing the nick must move from inside to outside the tandem gene array. If the repeat unit lacks an ARS, BITREx depends on replication initiated from an external ARS flanking the tandem gene array on the opposite side of the induced nick. As the array expands, the distance between the ARS and the nick increases, thus decreasing the likelihood that the outward replication fork will reach the nick earlier than the inward fork initiated from the nearest ARS on the same side as the nick. Consequently, the success rate of BITREx per cell cycle declines, limiting the maximum expansion range. In contrast, if the repeat unit contains an ARS, cells can more reliably maintain the desired replication fork direction. The internal ARS makes BITREx more autonomous and less dependent on an external ARS. Therefore, including an ARS in the repeat unit of a synthetic array is advantageous. Although autonomous BITREx theoretically allows for unlimited expansion, the increased instability of highly extended arrays realistically limits the practical extent of expansion.

Interestingly, BITREx requires nCas9 to nick the template DNA for leading strand synthesis, but not for lagging strand synthesis. This strand specificity likely reflects the asymmetry observed in the repair of replication-coupled DNA breaks, as revealed in recent studies on mammalian cells:^32,33^ nicks on the leading and lagging strand templates lead to the formation of seDSBs and deDSBs, respectively. The replisome notably bypasses nicks on the lagging strand templates to generate deDSBs directly, or without contribution from the converging replication fork, thereby likely preventing BIR. Intriguingly, one yeast study showed that a nick on a leading strand template induces a seDSB, while another nick on a different leading strand template induces a deDSB, independently of the converging replication fork.^34^ Although the determinants of these differential fates remain elusive, these variations may partly explain why ten out of the 16 gRNAs designed for lead collapse were ineffective.

While BITREx increased the average copy number of repeat units in the population, nanopore sequencing revealed the presence of contracted arrays. How does this contraction occur? If extensive 5′-to-3′ end-resection converts not only the terminal but also the second terminal repeat unit to ssDNA, the latter ssDNA may hybridize with the terminal repeat unit on the sister chromatid. In this scenario, the terminal repeat unit of the invading ssDNA strand is left behind as a flap. If this flap is degraded by flap endonucleases such as Rad1-Rad10, the invading ssDNA (i.e., the second terminal repeat unit) can initiate BIR, leading to the contraction of the tandem array. A similar situation could occur if the 3′-to-5′ exonuclease activity of Pol δ excessively degrades or “chews back” the invading strand.^35^ Therefore, appropriate suppression of these end resection activities may prevent contraction events and improve BITREx. These situations are more likely to occur when the repeat unit is short, meaning BITREx may not be effective in expanding micro- and mini-satellite DNAs, particularly when the repeat number is low.

We have successfully applied BITREx to an interrupted two-unit array containing four fluorescent protein genes as the intervening sequence, resulting in their overexpression to exert a dosage effect. These results suggest that when applying to a two-unit array with a biosynthetic gene cluster as the intervening sequence, BITREx can enhance the yield of the biosynthetic pathway’s product. In contrast to the serial configuration within a gene cluster, individual genes encoding pathway components can be distributed across multiple loci, where BITREx could act in parallel to increase their copy numbers. BITREx occurs stochastically in each cell cycle. Once it occurs, the subsequent cell division becomes asymmetric regarding unit copy number: one daughter cell inherits the donor chromatid with the original array, while the other inherits the acceptor chromatid with the expanded array. Additionally, the efficiency of BITREx varies from one locus to another. Therefore, parallel BITREx would create a cell population with diverse stoichiometry among pathway components, which could help identify an optimal pathway design for maximizing the yield of the pathway’s product.

It should be noted that BIR is less accurate than normal S-phase replication.^36^ During BIR, the Pif1 helicase immediately dissociates the newly synthesized leading strand DNA from its template, leaving it single-stranded until lagging strand synthesis occurs. As a result, the mismatch repair system functions ineffectively in BIR. Furthermore, the exposed nucleobases in ssDNA are much more susceptible to damage compared to those in dsDNA. Indeed, inducing BIR in yeast in the presence of an alkylating agent has led to the formation of mutation clusters similar to those found in cancer genomes.^37^ Therefore, we hypothesize that BITREx in the presence of mutagens, or mutagenic BITREx, will not only expand a tandem gene array but also diversify its repeat unit sequence, potentially generating an array of paralogs similar to OR gene loci.

We anticipate that BITREx will enable these and other unique applications in genome engineering.

### Limitations of the study

BITREx is a replication-coupled process and thus not applicable to non-dividing cells. By its nature, it cannot effectively expand tandem gene arrays located between actively firing nearby replication origins. Currently, predicting the performance of BITREx is challenging due to its reliance on various factors, including gRNA efficacy in inducing seDSB, the local environment of the target locus, and the composition of the repeat unit sequence, as some sequences can hinder BIR.^38^ Additionally, there is no method for stably maintaining tandem gene arrays that have been highly expanded by BITREx. Further investigation is needed to optimize BITREx for use in mammalian cells.

## ACKNOWLEDGEMENTS

We thank Tetsuya Hayashi and Yasuhiro Gotoh for the PFGE equipment. We appreciate the technical assistance from the Research Support Center of the Research Center for Human Disease Modeling at Kyushu University Graduate School of Medical Sciences, which is partially supported by the Mitsuaki Shiraishi Fund for Basic Medical Research. This work was funded by JST CREST Grant Number JPMJCR19S1.

## AUTHOR CONTRIBUTIONS

Conceptualization, H.T., S.O., and T.I.; Methodology, H.T., S.O., and G.D.; Investigation, H.T. and S.O.; Writing – Original Draft, H.T.; Writing – Review & Editing, H.T., S.O. and T.I.; Funding Acquisition, T.I.; Resources, H.T., S.O., G.D., Y.S., and E.K.; Supervision, T.I.

## DECLARATION OF GENERATIVE AI AND AI-ASSISTED TECHNOLOGIES

During the preparation of this work, the authors used ChatGPT to improve the readability of certain sentences. After using this tool/service, the authors reviewed and edited the content as needed and take full responsibility for the content of the publication.

## METHODS

### Yeast strains

All yeast strains used in this study are derived from BY4741 (*MAT***a** *his3*Δ1 *leu2*Δ0 *met15*Δ0 *ura3*Δ0)^39^ (Table S1). This study used standard culture media and genetic methods.^40^ We deleted a gene of interest by transforming yeast cells with a DNA fragment composed of a *KanMX* cassette sandwiched by the 5′- and 3′-flanking sequences of the open reading frame of the gene, which was amplified from the corresponding deletant strain in Yeast Deletion Clones *MAT***a** Complete Set (Invitrogen) using PCR primers listed in Table S2.

### Yeast plasmids

All plasmids used in this study are listed in Table S3. All primers for plasmid construction were purchased from Sigma-Aldrich and Eurofins Genomics. Plasmids were constructed by seamless cloning using HiFi DNA Assembly (NEB) or Golden Gate Assembly (NEB).

The integrative plasmid YIplac128-pGAL1-nCas9 (Cas9^D10A^ or Cas9^H840A/N854A^)-tADH1 (*LEU2*) harbors a gene encoding nCas9 derived from *Streptococcus pyogenes* fused with the SV40 nuclear localization signal as described previously^41^ under the control of the *GAL1* promoter. It was used for yeast transformation after AgeI digestion to be integrated into the *GAL1* promoter on the genome.

The integrative plasmid pFA6a-pCUP2-yGEV-tADH1-HphMX (HygR) harbors a gene encoding β-estradiol-responsive artificial transcription activator GEV^14^ under the control of the *CUP2* promoter. It was used for yeast transformation after MfeI digestion to be integrated into the *CUP2* promoter on the genome.

Centromeric plasmids for gRNA expression harbor a gRNA gene under the control of the *GAL1* promoter. The gRNA scaffold sequence contains a base-flip and a stem-loop extension for stable gRNA expression^42^. To cut off an unnecessary sequence from the 5′-terminal portion of the gRNA-containing transcript, each gRNA sequence is preceded by a hammerhead ribozyme (Table S3). To define the 3′-terminus, each gRNA sequence is followed by the HDV ribozyme on the *GAL1* promoter plasmid (Table S3). For designing gRNAs, CRISPRdirect^43^ was used to select target sites in the yeast genome listed in Table S4.

### Yeast genome editing

For constructing the *gRNA1inv*, *pif1-m2*, *rtt109-K290Q*, *cup1ru*Δ::*ymNG* array, *gRNA1ts-cup1ru*Δ::*NatMX*, *ho*Δ::2×*CUP1RU*, *ho*Δ::2×*ymNGRU, X-2*Δ::2×*CUP1RU*, *X-2*Δ::2×*ymNGRU* strains, we performed SpCas9 or enAsCas12a-based gene editing as described previously.^44^ All gene-editing plasmids used in this study are listed in Table S3.

### Yeast cell culture

Yeast cells were grown at 30°C overnight in 2 mL of SC−Ura, SC-His-Ura, SC−Leu−Ura, or SC−His−Leu−Ura medium supplemented with 2% glucose with or without G418 disulfate and/or hygromycin B (Nakalai tesque). On the following day, the OD_620_ of each sample was recorded, and 10–50 μL of the culture diluted up to 1 × 10^6^ times was inoculated into 2 or 5 mL of the fresh medium containing 10 nM β-estradiol, supplemented with or without 5 mM NAM. Genomic DNA was extracted from the remaining culture using the GC prep method for qPCR.^45^ The same process was repeated every 1 to 3 days. The division number per day was calculated from the change of OD_620_.

### Quantitative PCR (qPCR)

Genomic DNA was diluted ten times with distilled water before qPCR. Each qPCR solution (20 μL) contained 2 μL of diluted DNA, 10 μL of KOD SYBR qPCR Mix (TOYOBO), 0.04 μL of 50× ROX Reference Dye (TOYOBO), 2 pmol each of the forward and reverse primers. The primers used for qPCR are listed in Table S2. Each qPCR assay was performed in duplicate, using QuantStudio3 (Applied Biosystems) according to the manufacturer’s instructions. The amplification condition was initial denaturation at 98°C for 2 min followed by 40 times iteration of a 3-step thermal cycle composed of 98°C for 10 s, 55°C for 10 s, and 68°C for 30 s. All qPCR runs included 10-fold serial dilutions to generate standard curves. The quantity of *CUP1*, *ENA1, ymNG,* and *HIS3* was normalized to that of *ACT1*. The copy number of *CUP1*, *ENA1, ymNG,* and *HIS3* in the standard curves was calibrated by nanopore sequencing results in the BY4741 strain. The CNA/G for each gene was calculated with the below formula: CNA/G = (Copy number_Day T_ − Copy number_Day 0_) / Division number.

### Modeling the contraction of extended *CUP1* array

To interpret the plot between the initial copy number and CNA/G (Figure 4H), we deduced a theoretical plot assuming that contraction occurs via homologous recombination between two *CUP1* repeat units, following second-order kinetics.

Let *X* be the copy number of repeat units, and *k* be the rate constant of homologous recombination between the repeat units. The rate equation is given by:

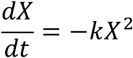

Let *X*_0_ be the initial repeat unit copy number. Then, solving this differential equation yields:

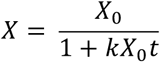

The copy number as a function of time decreases along a rectangular hyperbola. Since *CNA/G* is defined as the difference in copy number at *t* = *T* and *t* = 0 divided by the generation number *G*, it can be expressed as:

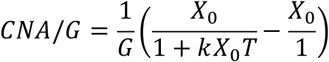

Simplifying this:

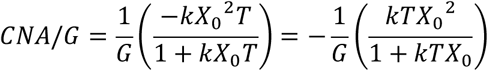

Finally:

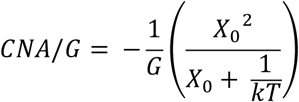

If *k* and *T* are constants and *X*_0_ is the variable, then *CNA*/*G* with respect to *X*_0_ decreases along the sum of a linear function and a rectangular hyperbola. As the initial copy number *X*_0_increases, the contraction rate *CNA*/*G* asymptotically approaches a straight line, as observed in Figure 4H.

### Nanopore sequencing

Genomic DNA was extracted using Monarch HMW DNA Extraction Kit for Tissue (NEB). We avoided vortexing to obtain high molecular weight DNA and used mixing by gentle pipetting with a wide-bore tip instead. DNA libraries were prepared using the ligation sequencing kit SQK-LSK109, SQK-LSK114 and the native barcoding kit EXP-NBD104, EXP-NBD114, or SQK-NBD114 (Oxford Nanopore Technologies) according to the manufacturer’s instructions. We modified the protocol of the ligation sequencing kit as follows: DNA fragmentation, omitted; duration of the enzymatic repair steps at 20 and 65°C, both extended from 5 min to 30 min; and the duration of the ligation step, extended from 10 to 30 min; incubation time for elusion with 0.4× AMPure XP, extended from 10 min to 20 min. The library was sequenced with the flowcell FLO-MIN106D R9.4.1 using the MinION sequencer and FLO-PRO114M R10.4.1 using the PromethION 2 Solo sequencer (Oxford Nanopore Technologies). MinKNOW software was used to control the MinION and PromethION devices. The run time was set to 72 h. Base calling was performed using Guppy v6.5.7 and Dorado v0.7.3. The assessment of sequencing data was performed using NanoPlot.^46^

### Dot plot analysis of nanopore reads

We used nanopore sequencing data in FASTA format to draw dot plots using YASS. ^47^ We first selected reads spanning the entire array using 1-kb upstream and 1-kb downstream sequences of the *CUP1*, *ENA1/2/5*, and *ymNeonGreen* (*ymNG*) array as queries of minialign (https://github.com/ocxtal/minialign) and then used these reads as the first input sequence for YASS. As the second input, we used the reference sequence of interest (*CUP1RU* for *CUP1* array and *ENA2RU* for *ENA1/2/5* array). By manually counting the diagonal lines in each dot plot, we determined the copy number of *CUP1RU* and *ENA2RU*.

### Translocation analysis using nanopore reads

We used nanopore sequencing data in FASTQ format, selected reads containing the *CUP1RU*, divided each read into 5′- and 3′-flanking regions of the *CUP1RU, and* extracted the flanking regions using Target_seq_extraction_single.py and Read_split_by_target.py (https://github.com/poccopen/Nanopore). We mapped reads to the S288c reference genome using Minimap2,^48^ SAMtools,^49^ and BEDtools.^50^ Data were visualized with the Integrative Genomics Viewer (IGV).^51^

### Copy number estimation from nanopore reads

We used nanopore sequencing data in FASTQ format and mapped reads to the S288c reference genome (version R64-2-1, http://sgd-archive.yeastgenome.org/sequence/S288C_reference/genome_releases/S288C_reference_genome_R64-2-1_20150113.tgz) using SAMtools^49^ and BEDtools,^50^ and then normalized read count of each nucleotide was calculated using Bedgraph_norm_ratio.py (https://doi.org/10.5281/zenodo.11515696). Data were visualized with the IGV.^51^

### Illumina sequencing

Genomic DNA was extracted using *Quick*-DNA Fungal/Bacterial Miniprep Kit (Zymo Research) and then fragmented to 300 bp using the S220 Focused-ultrasonicator (Covaris). DNA libraries with indexing were prepared using the ThruPLEX DNA-Seq kit (TaKaRa). We used the DNA Single Index Kit – 12S Set A or B (TaKaRa) for indexing according to the manufacturer’s instructions. The library was sequenced with MiSeq Reagent Kit v3 using the MiSeq instrument (Illumina). We mapped 2 × 75 bp reads to the S288c reference genome using Bowtie2, ^52^ SAMtools,^49^ and BEDtools,^50^ and then normalized read count of each nucleotide was calculated using Bedgraph_norm_ratio.py (https://doi.org/10.5281/zenodo.11515696). Data were visualized with the IGV.^51^

### Pulsed-field gel electrophoresis and Southern blot hybridization

Agarose-embedded yeast DNA was prepared using CHEF Genomic DNA Plug Kits (BioRad) according to the manufacturer’s instructions. DNA digested with or without EcoRI was subjected to 1% and 0.8% Certified Megabase Agarose (Bio-Rad) in 0.5× TBE and 1× TAE, respectively. Pulsed-field gel electrophoresis (PFGE) was performed using CHEF mapper XA (BioRad) according to the manufacturer’s instructions. PFGE running conditions were a 60 -120 s pulse time, 120° angle, and 6 V/cm for 24 h at 14°C in a 1% agarose gel and 500 s pulse time, 106° angle, and 3 V/cm for 48 h at 14°C in a 0.8% agarose gel. The gel was then stained with SYBR Green I Nucleic Acid Gel Stain (Invitrogen) at room temperature for 30 min with shaking, destained in distilled water for 1 h, and the fluorescence signals were detected with ChemiDocTouch system (Bio-Rad). Transfer to the membrane was performed using Hybond-N+ (Cytiva) according to the manufacturer’s instructions. The blot was hybridized with a *CUP1* probe or outside probes at 55°C overnight after UV-crosslinking. The probe was generated by PCR using the primers listed in Table S2, followed by labeling with alkaline phosphatase using the labeling module of the AlkPhos Direct Labelling and Detection System kit (Cytiva). Following appropriate blot washing, chemiluminescent signals were generated using the CDP-Star Detection Reagent in the kit and detected with the ChemiDocTouch system (Bio -Rad). Images were processed with ImageJ software (National Institutes of Health).

### Immunoblot analysis

The amount of histone H3 and acetylated histone H3 lysine 56 (H3K56ac) was analyzed by western blotting. Proteins were extracted as described previously,^53^ and separated with sodium dodecyl sulfate-polyacrylamide gel electrophoresis using 12% Mini-PROTEAN TGX Precast Protein Gel (Bio-Rad). Transfer to the membrane was performed with iBind Western System (Thermo Fisher Scientific) according to the manufacturer’s protocol. Primary antibodies to detect histone H3 and H3K56ac were Anti-Histone H3 Antibody, CT, pan, clone A3S, rabbit monoclonal antibody (1:500, Sigma-Aldrich), and Histone H3K56ac rabbit polyclonal antibody (1:2500, Active Motif), respectively. The secondary antibody was Anti-rabbit IgG, HRP-linked Antibody (1:2000, Cell Signaling Technology). Following incubation with Clarity Western ECL Substrate (Bio-Rad), chemiluminescent signals were detected with the ChemiDocTouch system (Bio -Rad). Gel images were processed with ImageJ software.

### Fluorescence microscopy and image processing for yeast cells

Image acquisitions of yeast cells were performed on a microscope (Ti-E, Nikon, Tokyo, Japan) with a 20× objective lens (CFI Plan Apo Lambda 20X, MRD00205, Nikon), a sCMOS camera (ORCA-Fusion BT, C15440-20UP, Hamamatsu photonics, Hamamatsu, Japan), and a solid-state illumination light source (SOLA SE II, Lumencor, Beaverton, OR, USA). Image acquisition was controlled by NIS-Elements version 5.3 (Nikon). Z-stacks were 7 × 0.9 μm. For imaging of mNeonGreen, a filter set (LED-YFP-A, Semrock, Rochester, NY, USA) was used with excitation light power set at 7% and the exposure time set at 200 msec/frame. For imaging of mCherry, a filter set (LED-TRITC-A, Semrock) was used with excitation light power set at 30% and exposure time set at 300 msec/frame. For imaging of miRFP682, a filter set (LED-Cy5.5-A, Semrock) was used with excitation light power set at 10% and exposure time set at 200 msec/frame. For imaging of TagBFP, a filter set (LED-DAPI-A, Semrock) was used with excitation light power set at 50% and exposure time set at 700 msec/frame. For DIC (differential interference contrast) image acquisition, the exposure time was set at 50 ms/frame. DIC images were captured only at the middle position of the Z-stacks.

Image processing and analysis were performed using Fiji.^54^ To generate 2-dimensional images of fluorescence channels from Z-stacks, background subtraction (sliding paraboloid radius set at 5 pixels with disabled smoothing) and maximum projection using 7 Z-slices were performed. Maximum projected fluorescence images and corresponding smoothed DIC images were superimposed. After global adjusting of brightness and contrast and cropping of the images, sequences of representative images were generated.

### HEK293T cell culture and transfection

Human embryonic kidney 293T cell (HEK293T) cells were cultured in Dulbecco’s modified Eagle’s medium (DMEM) (Gibco) supplemented with 10% fetal bovine serum (Gibco) and 1% penicillin–streptomycin (Gibco). Cells were maintained at 37°C in a humidified atmosphere containing 5% CO_2_. HEK293T cells were transfected with the FP-EGF-mCherry-PuroR plasmid using Lipofectamine 3000 reagent (Invitrogen) and FP-EGF-mCherry-PuroR was knocked-in at the *AAVS1* locus using the VIKING method.^29^ Puromycin (0.3 µg/µL) was added to the culture at 24 h after transfection. Following 48 h of cultivation, the cells were transfected with the nCas9-PuroR plasmid. Puromycin (2.0 µg/µL) was then added to the culture at 24 h after the second transfection, and the cells were grown for an additional 72 h.

### Flow cytometric analysis of transfected HEK293T cells

Transfected HEK293T cells were washed twice with phosphate-buffered saline (PBS) and detached from the dishes using 0.25% trypsin-EDTA (Gibco). The cells were then resuspended in PBS containing 0.2% bovine serum albumin (Thermo Fisher Scientific) and filtered through a 50 µm nylon mesh to obtain a single-cell suspension. Cell density was adjusted to 5 × 10^6^ cells/mL. Cells were analyzed using a BD FACSAria^TM^ Fusion cell sorter (BD Biosciences). Data acquisition was performed using BD FACSDiva^TM^ software (BD Biosciences). At least 10,000 events were collected per sample. Data were analyzed using FlowJo software (BD Biosciences), and gates were set based on isotype controls.

### Flow sorting of EGFP-positive HEK293T cells

EGFP and mCherry-positive cells were sorted using a BD FACSAria^TM^ Fusion cell sorter (BD Biosciences). The excitation wavelength for EGFP was set to 488 nm, and EGFP florescence was detected using a 530/30 nm bandpass filter. The excitation wavelength for mCherry was set to 561 nm, and mCherry fluorescence was detected using a 610/20 nm bandpass filter. Non-transfected HEK293T cells were used as negative controls to set the gates for GFP and mCherry-positive cells. At least 10,000 events were collected per sample.

### Fluorescence microscopy and image processing for HEK293T cells

Image acquisitions of HEK293T cells were performed on an imaging system (EVOS M7000, Thermo Fisher Scientific) with a 20× objective lens (NA 0.70, Olympus, AMEP4765EO). For imaging of EGFP, a filter set (EVOS Light Cube GFP 2.0, Thermo Fisher Scientific, AMEP4951) was used with the exposure time set at 2 msec/frame. For imaging of mCherry, a filter set (EVOS Light Cube Texas Red 2.0, Thermo Fisher Scientific, AMEP4955) was used with the exposure time set at 50 msec/frame. For bright field image acquisition, the exposure time was set at 10 ms/frame. Image processing and analysis were performed using Fiji.^53^ Background subtraction (sliding paraboloid radius set at 5 pixels with disabled smoothing) were performed. Fluorescence images and corresponding bright field images were superimposed. After global adjusting of brightness and contrast and cropping of the images, sequences of representative images were generated.

### Genomic DNA preparation from HEK293T cells

Genomic DNA was extracted from HEK293T cells using NucleoSpin Tissue (Macherey-Nagel) and Monarch HMW DNA Extraction Kit for Cells & Blood (Monarch) according to the manufacturer’s protocol. Extracted DNA was used for preparation of library DNA for Nanopore sequencing as described above.

## QUANTIFICATION AND STATISTICAL ANALYSIS

One-way ANOVA test and Student’s t-test were employed to calculate p values, as indicated in the figure legends. In general, results were considered statistically significant when p < 0.05.

## DATA AVAILABILITY

All raw sequencing data used in this study were deposited with links to BioProject accession numbers PRJBD18647, PRJBD18687, and PRJBD18705 in the DDBJ BioProject database.

## SUPPLEMENTAL INFORMATION

Table S1. Yeast strains used in this study.

Table S2. PCR primers used in this study.

Table S3. Plasmids used in this study.

Table S4. gRNAs and crRNAs used in this study.

