## Supplemental Data 1 for "Strategic targeting of Cas9 nickase expands tandem gene arrays"

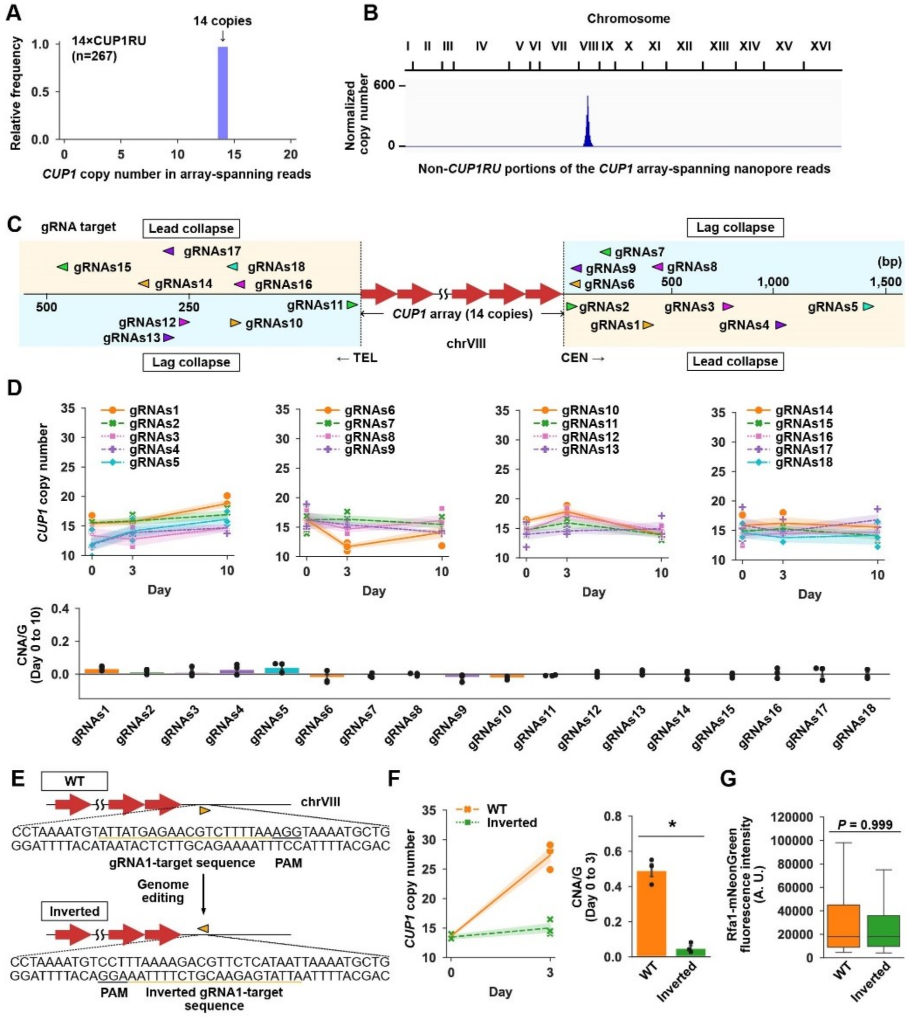


**Figure S1: BITREx of *CUP1* array, related to Figure 1**

(A) *CUP1* array length of the parental strain used in this study. Nanopore reads containing both the 5′- and 3′-flanking regions of *CUP1* array were used to determine the distribution of *CUP1RU* copy number.

(B) Genomic location of *CUP1* array. Nanopore reads containing *CUP1RU* were selected and their non-*CUP1RU* portions were mapped to the reference genome sequence.

(C) *CUP1* array and target sites of ineffective gRNAs (gRNAs1–gRNAs18). Similar to Figure 1C.

(D) Performance of 18 gRNAs shown in (C). Similar to Figures 1D and 1E. Shading and error bar, SD (n = 3 biological replicates).

(E) Inversion of the gRNA1 target sequence.

(F) BITREx in the wild-type and inverted strains. Left, CNA of *CUP1*; right. Shading and error bar, SD (n = 3 biological replicates).

(G) Box plots showing Rfa1-mNeonGreen fluorescence in wild-type and inverted strains with Cas9 and gRNA1. Since Rfa1 accumulates on ssDNA generated by end-resection at DSB sites as a component of the RPA complex, the fluorescence intensity serves as an indicator of gRNA1-guided Cas9 cleavage.

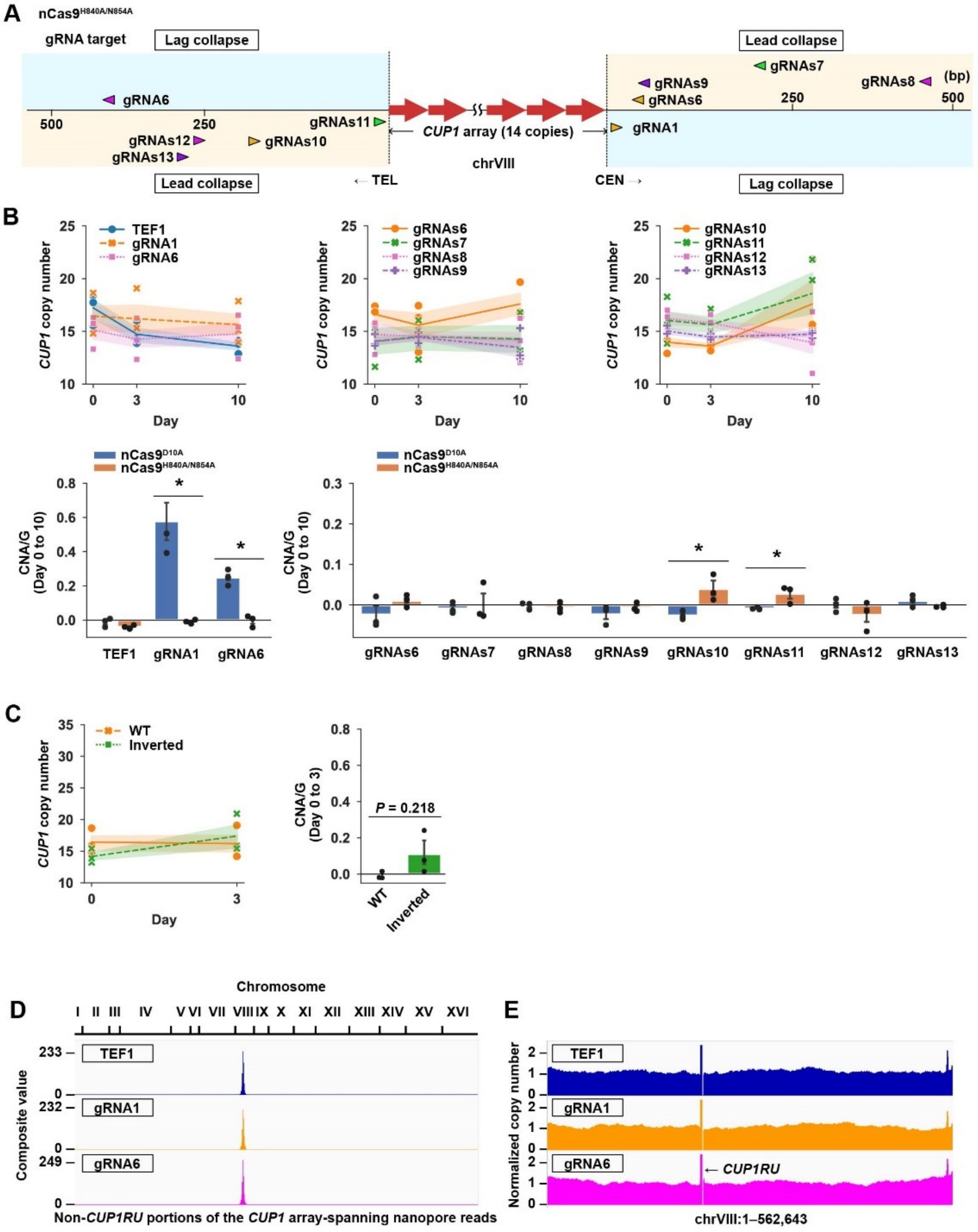


*(legend on next page)*

**Figure S2: Requirement of lead collapse for BITREx, related to Figure 1**

(A) *CUP1* array and target sites of 10 gRNAs used with nCas9$^{H840A/N854A}$. Two gRNAs (gRNA1 and gRNA6) were effective when used with nCas9$^{D10A}$ and should induce lag collapse when used with nCas9$^{H840A/N854A}$. The remaining eight gRNAs (gRNAs6–gRNAs13) were ineffective when used with nCas9$^{D10A}$ (Figure S1D) and should induce lead collapse when used with nCas9$^{H840A/N854A}$.

(B) BITREx using nCas9$^{H840A/N854A}$. Similar to Figures 1D and 1E. Shading and error bar, SD (n = 3 biological replicates). *P < 0.05 (Student's *t*-test)

(C) BITREx using gRNA1 and nCas9$^{H840A/N854A}$ in the inverted strain (Figure S1E). Shading and error bar, SD (n = 3 biological replicates).

(D) Genomic location of *CUP1* array. Nanopore reads containing *CUP1RU* were selected and their non-*CUP1RU* portions were mapped to the reference genome sequence.

(E) Normalized read counts across chromosome VIII in nanopore sequencing. Read counts were normalized to the average counts of genomic regions excluding rRNA, *CUP1RU*, Ty elements, and mitochondrial DNA. Note that while the sacCer3 reference genome sequence contains two copies of *CUP1RU*, the second copy is masked with 'N' prior to mapping. Consequently, the normalized read count directly reflects the *CUP1RU* copy number. The gap in read counts adjacent to *CUP1RU* is due to the aforementioned masking.

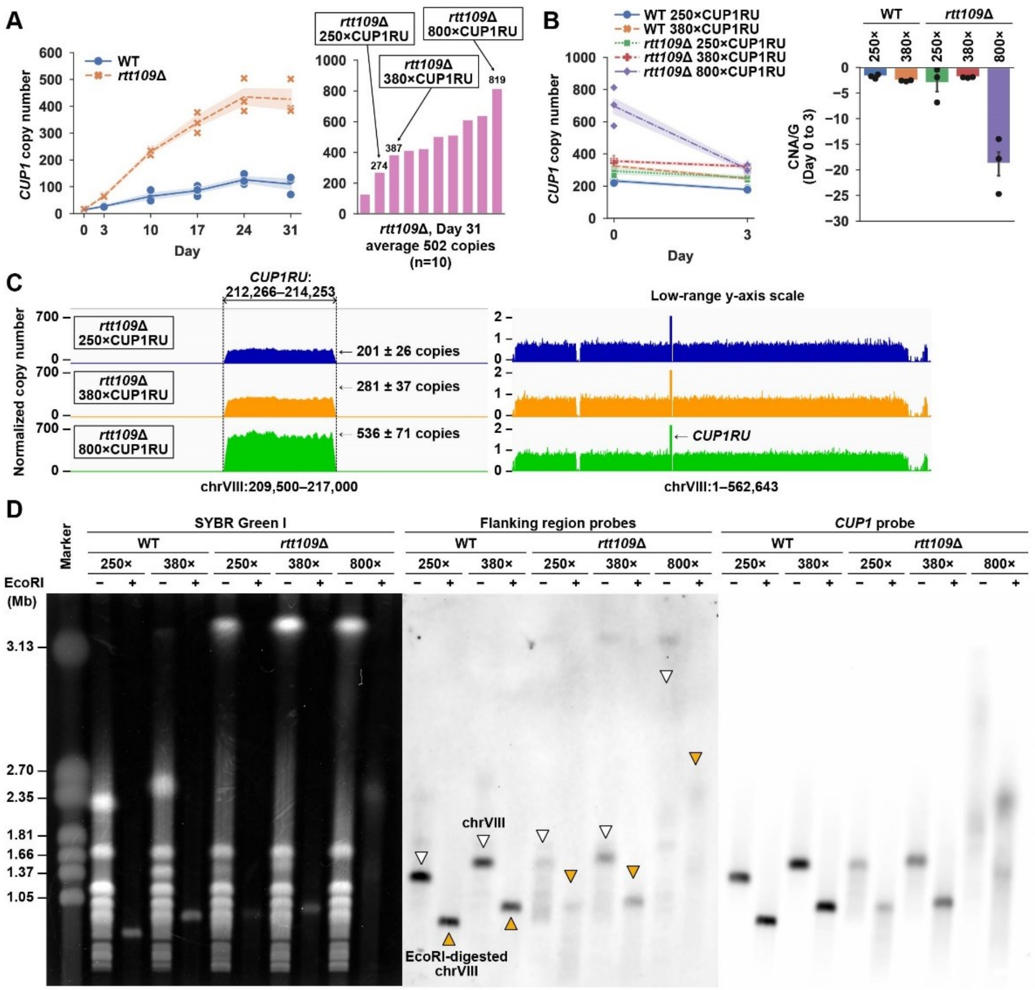


**Figure S3: Long-term BITREx in the absence of Rtt109, related to Figure 3**

(A) CNA of *CUP1* over the 31-day BITREx with gRNA1 in *rtt109*Δ cells. Shading, SD (n = 3 biological replicates). *CUP1* copy numbers of 10 randomly picked clones on day 31 are shown in the right panel.

(B) Stability of extended *CUP1* arrays. Each strain was cultivated for 3 days without BITREx induction. Shading and error bar, SD (n = 3 biological replicates).

(C) Normalized read counts in Illumina sequencing. Similar to Figure 3F.

(D) PFGE analysis of *CUP1* arrays expanded by long-term BITREx. Similar to Figure 3H. Note that most slowly migrating bands are chromosome XII, which carries the rDNA array showing remarkable expansion in *rtt109*Δ cells.

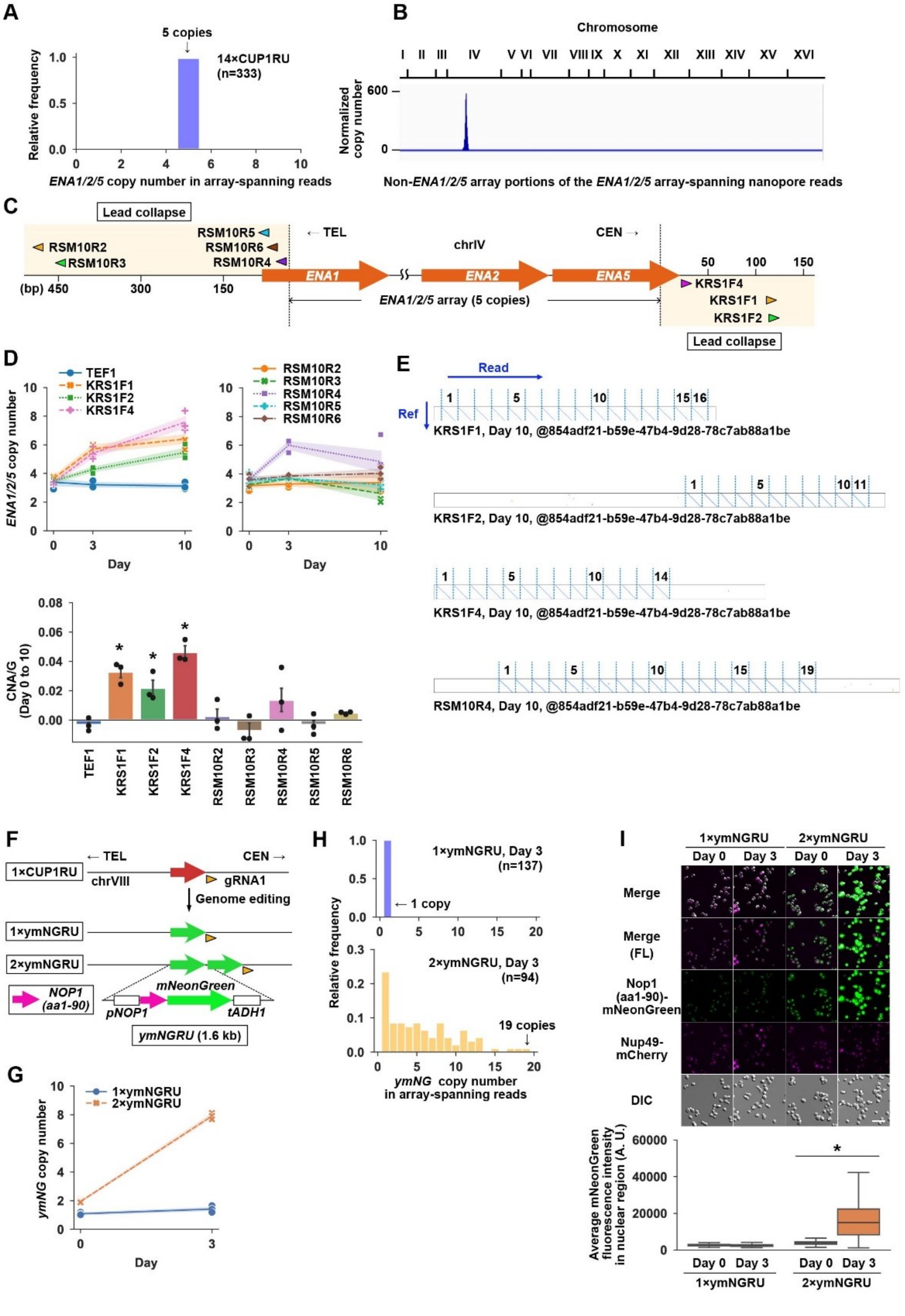


*(legend on next page)*

**Figure S4: BITREx of non-*CUP1* array, related to Figure 5**

(A) *ENA1/2/5* array length of the parental strain used in this study. Nanopore reads containing both the 5′- and 3′-flanking regions of *ENA1/2/5* array were used to determine the distribution of *CUP1RU* copy number.

(B) Genomic location of *ENA1/2/5* array. Nanopore reads containing *ENA1/2/5* were selected and their non-*ENA1/2/5* portions were mapped to the reference genome sequence.

(C) *ENA1/2/5* array and target sites of gRNAs tested in this study. Similar to Figure 1C.

(D) Performance of eight gRNAs shown in (C). Similar to Figures 1D and 1E. Shading and error bar, SD (n = 3 biological replicates).

(E) Representative dot plots comparing nanopore reads to the reference sequence of *ENA1/2/5* repeat unit. Genomic DNAs prepared from the cells with four gRNAs (KRS1F1, KRS1F2, KRS1F4, or RSM10R4) on day 10 were used for the nanopore sequencing.

(F) *ymNGRU* arrays generated on chromosome VIII. A single *CUP1RU* at the *CUP1* locus on chromosome VIII was replaced by a single copy or tandemly duplicated copies of *ymNGRU* using genome editing. *pNOP1*, *NOP1* promoter; *tADH1*, *ADH1* terminator.

(G) CNA of *ymNG* by BITREx with gRNA 1 in the strain bearing either a single copy *ymNGRU* (1×ymNGRU) or a two-unit *ymNGRU* array (2×ymNGRU). Shading, SD (n = 3 biological replicates).

(H) Distribution of *ymNGRU* copy number in nanopore reads spanning the entire array in the 1×ymNGRU and 2×ymNGRU strains on day 3 of BITREx.

(I) Fluorescence microscopic analysis of 1×ymNGRU and 2×ymNGRU strains. Upper panel, representative microscopic images. These strains have *NUP49-mCherry* to visualize the nuclei (magenta). FL, fluorescence; DIC, differential interference contrast. Lower panel, quantification of fluorescence intensity. Box plots indicate the distribution of the average mNeonGreen fluorescence intensity in the nuclear region. The bottom and top of the box show the first and third quartiles, respectively. The bar in each box represents the median value, and the error bars represent the range of values. *P < 0.001 (one-way ANOVA test).

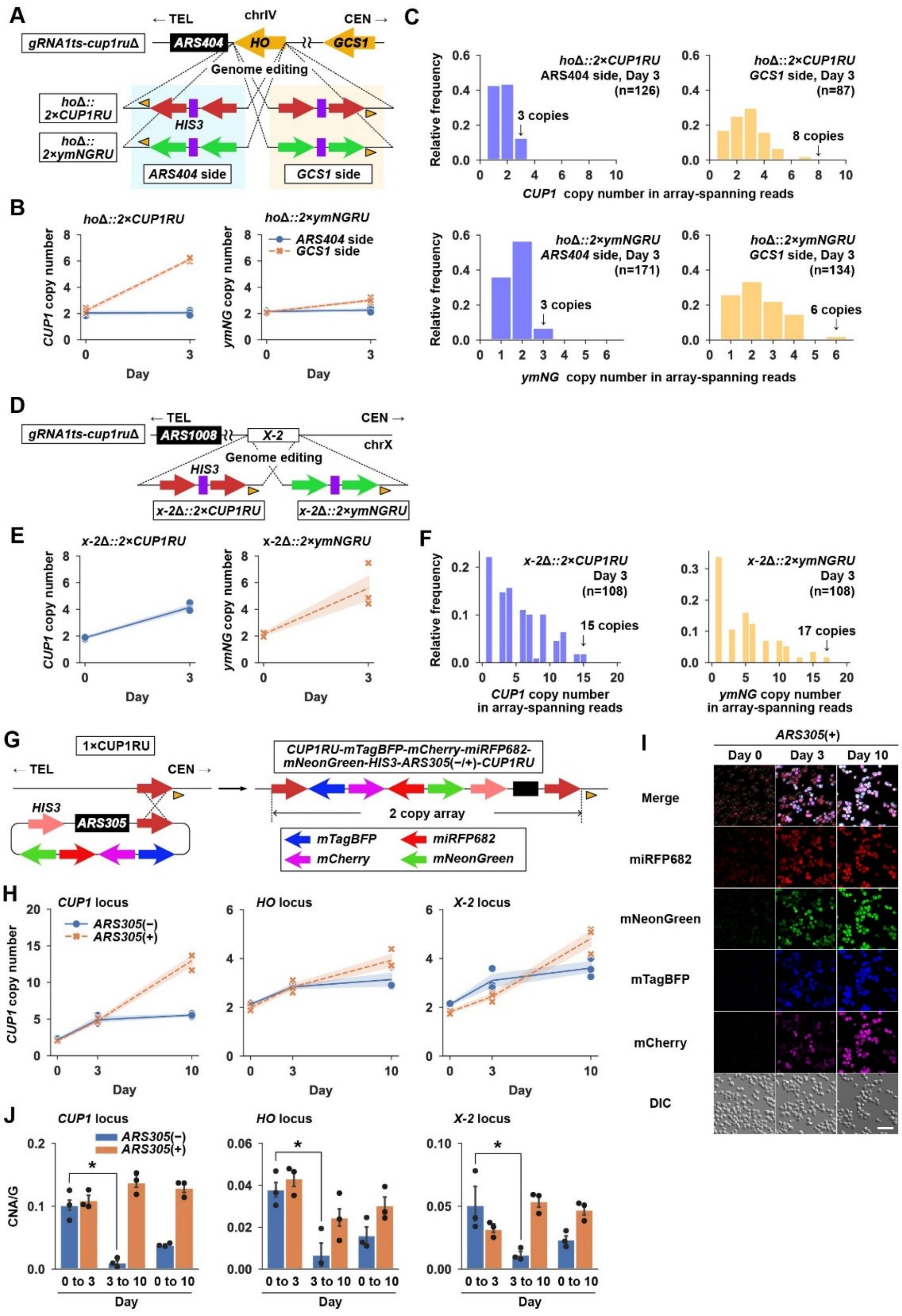


(legend on next page)

**Figure S5: BITREx of interrupted two-unit arrays, related to Figure 5**

(A) Interrupted two-unit arrays of *CUP1RU* and *ymNGRU* inserted at the *HO* locus on chromosome IV. Note that the two-unit arrays were inserted in two orientations. Orange arrowheads, gRNA1-target site. Note that the *HO* locus is flanked by *ARS404* and *GCS1*.

(B) CNA of *CUP1* and *ymNG* in the *ho*Δ*::2×CUP1RU* and *ho*Δ*::2×ymNGRU* strains, respectively. Shading, SD (n = 3 biological replicates).

(C) Distribution of *CUP1/ymNG* copy number in nanopore reads spanning the entire array obtained on day 3.

(D) Interrupted two-unit arrays of *CUP1RU* and *ymNGRU* integrated to the *X-2* locus on chromosome X. *ARS1008* is the nearest ARS in the side opposite to the gRNA1-target site (orange arrowhead).

(E) CNA of *CUP1* and *ymNG* in the *x-2::2×CUP1RU* and *x-2::2×ymNGRU* strains, respectively. Shading, SD (n = 3 biological replicates).

(F) Distribution of *CUP1/ymNG* copy number in nanopore reads spanning the entire array obtained on day 3.

(G) Two-unit *CUP1* array interrupted by an intervening sequence bearing four fluorescent protein genes. The array was generated by the recombination of a plasmid bearing a *CUP1RU* and the intervening sequence with the single-copy *CUP1RU*.

(H) CNA of *CUP1* in the strains bearing the interrupted *CUP1* array in (G) at *CUP1*, *HO*, or X-2 loci. Isogenic stains without the embedded *ARS305* were also shown. Shading, SD (n = 3 biological replicates). *P < 0.05 (Student's *t*-test).

(I) Microscopic images of the strain bearing the interrupted *CUP1* array in (G). Cells were subjected to fluorescence microscopy on days 0, 3, and 10. Red, miRFP682; green, mNeonGreen; blue, mTagBFP; magenta, mCherry; grey, DIC.

(J) Alteration of CNA/G during BITREx. CNA/G was calculated for the periods from day 0 to 3, day 3 to 10, and day 0 to 10. *P < 0.05 (Student's *t*-test).

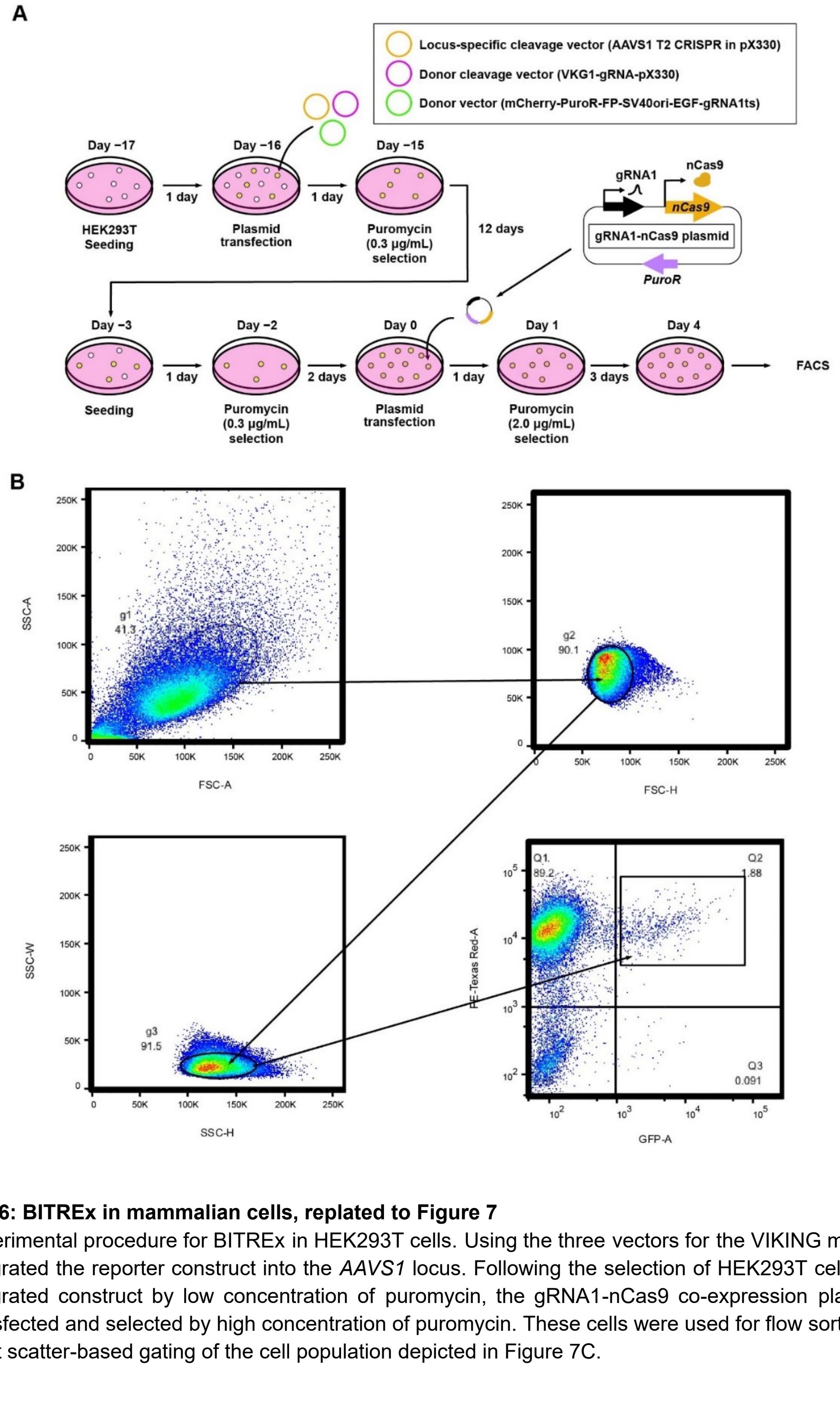


**Figure S6: BITREx in mammalian cells, replated to Figure 7**

(A) Experimental procedure for BITREx in HEK293T cells. Using the three vectors for the VIKING method, we integrated the reporter construct into the *AAVS1* locus. Following the selection of HEK293T cells with the integrated construct by low concentration of puromycin, the gRNA1-nCas9 co-expression plasmid was transfected and selected by high concentration of puromycin. These cells were used for flow sorting.

(B) Light scatter-based gating of the cell population depicted in Figure 7C.
